# A nucleotide resolution map of Top2-linked DNA breaks in the yeast and human genome

**DOI:** 10.1101/530667

**Authors:** William Gittens, Dominic J. Johnson, Rachal M. Allison, Tim J. Cooper, Holly Thomas, Matthew J Neale

**Affiliations:** Genome Damage and Stability Centre, University of Sussex, UK, BN1 9RQ

## Abstract

DNA topoisomerases are required to resolve DNA topological stress. Despite this essential role, abortive topoisomerase activity generates aberrant protein-linked DNA breaks, jeopardising genome stability. Here, to understand the genomic distribution and mechanisms underpinning topoisomerase-induced DNA breaks, we map Top2 DNA cleavage with strand-specific nucleotide resolution across the *S. cerevisiae* and human genomes—and use the meiotic Spo11 protein to validate the broad applicability of this method to explore the role of diverse topoisomerase family members. Our data characterises Mre11-dependent repair in yeast, and defines two strikingly different fractions of Top2 activity in humans: tightly localised CTCF-proximal, and broadly distributed transcription-proximal, the latter correlated with gene length and expression. Moreover, single nucleotide accuracy enables us to reveal the influence primary DNA sequence has upon Top2 cleavage—distinguishing canonical DNA double-strand breaks (DSBs) from a major population of DNA single-strand breaks (SSBs) induced by etoposide (VP16) *in vivo*.

## Introduction

DNA topoisomerases are a broad and ubiquitous family of enzymes that tackle topological constraints to replication, transcription, the maintenance of genome structure and chromosome segregation in mitosis and meiosis (Pommier et al., 2016). Although the specific mechanisms by which this is accomplished vary considerably across the family, key aspects are shared: including single or double-strand DNA cleavage to form a transient covalent complex (CC), which allows alteration of the topology of the nucleic acid substrate prior to religation (Wang, 2009). These processes are essential but carry with them a significant risk to genome stability because the CC may be stabilised as a permanent protein-linked DNA break by several physiological factors; such as the proximity of other DNA lesions, the collision of transcription and replication complexes, denaturation of the topoisomerase, or by the binding of small molecules that inhibit religation (Reviewed in (Pommier et al., 2016; Nitiss, 2009)). Topoisomerase-induced DNA strand breaks have been proposed to constitute a significant fraction of the total damage to genomic DNA per day, and have been linked to the genesis and development of various cancers (Lin et al., 2009; Haffner et al., 2010), including a subset of therapy-related acute myeloid leukaemias (t-AML) caused by the use of Topoisomerase 2 (Top2) poisons in the chemotherapeutic treatment of primary cancers (Rowley and Olney, 2002; Cowell et al., 2012).

In *S. cerevisiae*, either Top1 or Top2 activity is sufficient to support transcription (Brill et al., 1987). However, whilst *top1*Δ cells are viable, Top2 is essential for sister chromatid segregation (Baxter and Diffley, 2008; DiNardo et al., 1984; Holm et al., 1985). In contrast to *S. cerevisiae*, all known vertebrate species encode two Top2 proteins (TOP2α and TOP2β). Interestingly, whilst TOP2α is essential for cellular proliferation—cells arrest in mitosis in its absence (Akimitsu et al., 2003)—TOP2β is not. Rather, TOP2β apparently plays an important role in promoting transcriptional programmes associated with neuronal development (Tiwari et al., 2012; Gómez-Herreros et al., 2014), a function that cannot be supported by TOP2α.

Accurate maps of the positions of topoisomerase-DNA covalent complexes (Top2 CCs) genome-wide and throughout the cell cycle provide insights into topological genome structural organisation, as well as providing the tools to study their repair should they become permanent DNA lesions. Whilst the dual catalytic sites present within the homodimeric Top2 enzyme suggest a primary role in DNA double-strand break (DSB) formation driven by a biological need for decatenation (Pommier et al., 2016), previous research has suggested that etoposide and other Top2 poisons may induce a population of DNA single-strand breaks (SSBs), due to independent inhibition of each active site (Wozniak and Ross, 1983; Long et al., 1986; Bromberg et al., 2003). Yet, despite the use of etoposide in chemotherapy, and the different toxicity of these two classes of DNA lesion, the prevalence of Top2-SSBs remains unclear.

More recently, general DSB mapping techniques have been applied to map etoposide-induced lesions (Canela et al., 2017; Yan et al., 2017). However, such methods are only able to map DSB ends, which therefore excludes and obscures etoposide-induced SSBs. Moreover, these methodologies lack specificity for protein-linked DNA ends, and require nucleolytic processing to blunt 5’ DNA termini as part of sample preparation (Canela et al., 2017). When combined, these factors lead to loss of nucleotide resolution at the site of Top2 cleavage.

To generate a more complete, and direct, picture of topoisomerase action across the genome, we present here a technique for nucleotide resolution mapping of protein-linked SSBs and DSBs (referred to collectively here as “Covalent Complexes”; CC), which we demonstrate to be generally applicable in both yeast and human cell systems. We establish with high confidence that the technique is able to map both the positions of Top2 CCs and also CCs of DNA linked to the meiotic recombination protein Spo11, which is related to the archaeal Topoisomerase VI (Bergerat et al., 1997). Furthermore, we provide insights into the spatial distribution of human TOP2 CCs, including comparative analysis of their local enrichment around transcription start sites (TSSs) and CTCF-binding motifs. We find that TSS-proximal TOP2 CC levels are strongly correlated with transcription, a question that has come under scrutiny recently (Canela et al., 2017). Finally, we present compelling evidence that etoposide induces a majority of Top2-linked SSBs *in vivo*, corroborating previous research and revealing, for the first time, that this is governed by primary DNA sequence at the cleavage site.

## Results

### CC-seq enriches Spo11-linked DNA fragments

To elucidate the *in vivo* functions of topoisomerase-like enzymes, we set out to establish a direct method (termed ‘CC-seq’) to enrich and map protein-DNA covalent complexes (CC) genome-wide with nucleotide resolution. Silica fibre-based enrichment of CCs has been used in the original identification of the meiotic DSB-inducer Spo11 (Keeney and Kleckner, 1995; Keeney et al., 1997). We first verified this enrichment principle using meiotic *sae2*Δ cells, in which Spo11-linked DSBs accumulate at defined loci due to abrogation of the nucleolytic pathway that releases Spo11 (Keeney and Kleckner, 1995). To demonstrate specific enrichment of protein-linked molecules, genomic DNA from meiotic *S. cerevisiae* cells was isolated in the absence of proteolysis, digested with *Pst*I restriction enzyme, and isolated on glass-fibre spin columns (**Figure 1A, Methods**). We used eluted material to assay a known Spo11-DSB hotspot by Southern blotting (**Figure 1B**). While DSB fragments are a minor fraction of input material (∼10% of total), and were absent in wash fractions, DSBs accounted for >99% of eluted material, indicating ∼1000-fold enrichment relative to non-protein-linked DNA (**Figure 1B**).

**Figure 1.**
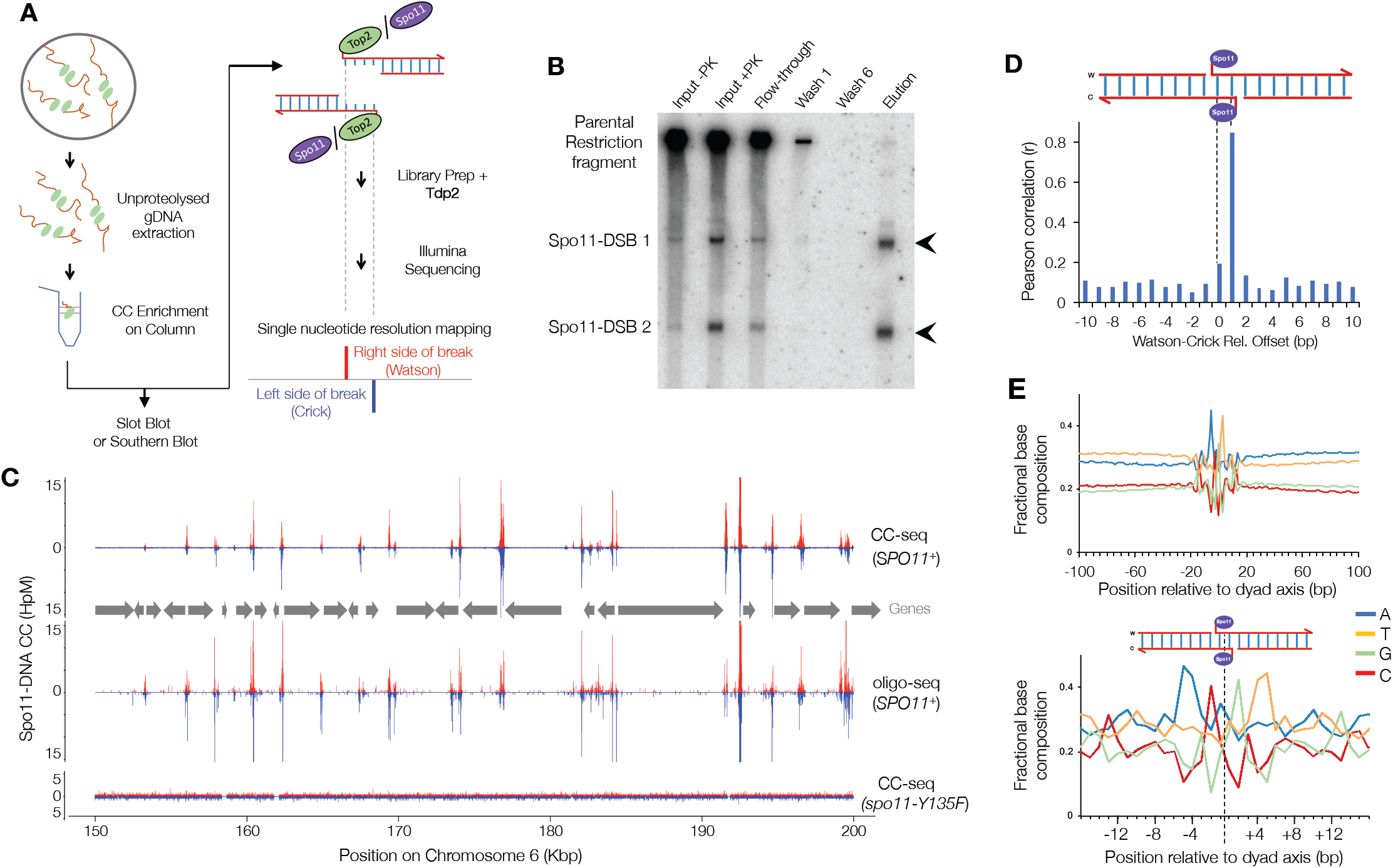
CC-seq maps covalent Spo11-linked DNA breaks in *S. cerevisiae* meiosis with nucleotide accuracy. **A)** Schematic of the CC-seq method. **B)** Column-based enrichment of Spo11-linked DNA fragments detected by Southern blotting at the *his4*∷*LEU2* recombination hotspot (**Methods**). Arrows indicate expected sizes of Spo11-DSBs. **C)** Nucleotide resolution mapping of *S. cerevisiae* Spo11 hotspots by CC-seq or oligo-seq (Pan et al., 2011). Red and blue traces indicate Spo11-linked 5’ DNA termini on the Watson and Crick strands, respectively. Grey arrows indicate positions of gene open reading frames. **D)** Pearson correlation (r) of Spo11 CC-seq signal between Watson and Crick strands, offset by the indicated distances. **E)** Average nucleotide composition over a 200 (top) and 30 bp (bottom) window centred on Spo11 breaks. Bases reported are for the top strand only. HpM = Hits per million mapped reads per base pair.

### CC-seq maps known Spo11-DSB hotspots genome-wide with high reproducibility

To generate a genome-wide map of Spo11-DSBs, genomic DNA from meiotic *sae2*Δ cells was sonicated to <400 bp in length, enriched upon the silica column, eluted, and ligated to DNA adapters in a two-step procedure that utilised the known phosphotyrosyl-unlinking activity of mammalian TDP2 to uncap the Spo11-bound end (**Figure 1A / Methods**) (Cortes Ledesma et al., 2009; Johnson et al., 2019). Libraries were paired-end sequenced and mapped to the *S. cerevisiae* reference genome (**Table S1**) alongside reads from a previous mapping technique (‘Spo11-oligo-seq’) that relies on the isolation of Spo11-linked oligonucleotides generated in wild-type cells during DSB repair (Pan et al., 2011). CC-seq revealed sharp, localised peaks (‘hotspots’) in *SPO11*+ cells that visually (**Figure 1C**) and quantitatively (**Figure S1A**, r=0.82) correlate with Spo11-oligo seq, and that are absent in the *spo11-Y135F* control strain in which Spo11-DSBs do not form (Bergerat et al., 1997), demonstrating that signal detected by CC-seq accurately reflects the distribution of Spo11 cleavages. Moreover, CC-seq biological replicates were highly correlated (**Figure S1B**; r = 0.98), demonstrating high reproducibility. Finally, when analysed at nucleotide resolution, CC-seq revealed high correlation (r = 0.85) between the frequency of Watson-mapping and Crick-mapping cleavage sites, when offset by a single bp (**Figure 1D and S1C**). This correlation is expected because of the 2 bp 5’ overhang generated at Spo11-DSBs *in vivo* (**Fig1D**; e.g. (Liu et al., 1995))—demonstrating the nucleotide resolution accuracy of CC-seq, which is further supported by our observation of nucleotide composition preference at the site of cleavage (**Figure 1E**).

### CC-seq enriches and maps Top2-linked DNA fragments from *S. cerevisiae*

Confident in the specificity of CC-seq to map meiotic protein-linked DSBs with high resolution and dynamic range, we employed CC-seq to characterise within *S. cerevisiae* the covalent complexes generated naturally by Topoisomerase 2 (Top2) that become stabilised upon exposure to etoposide, and are thus a proxy for Top2 catalytic activity (Minocha and Long, 1984; Chen et al., 1984). Because *S. cerevisiae* is relatively insensitive to etoposide, we utilised strains (‘*pdr1*Δ’) in which the action of major drug export pathways are downregulated due to modulation of *PDR1* activity (Stepanov et al., 2008). As expected, *pdr1*Δ dramatically increased cellular sensitivity to etoposide, which was further enhanced by mutation of *SAE2* and *MRE11* (**Figure 2A**), factors involved in the repair of covalent protein-linked DNA breaks (Neale et al., 2005; Cannavo et al., 2018; Hoa et al., 2016; Hartsuiker et al., 2009).

**Figure 2.**
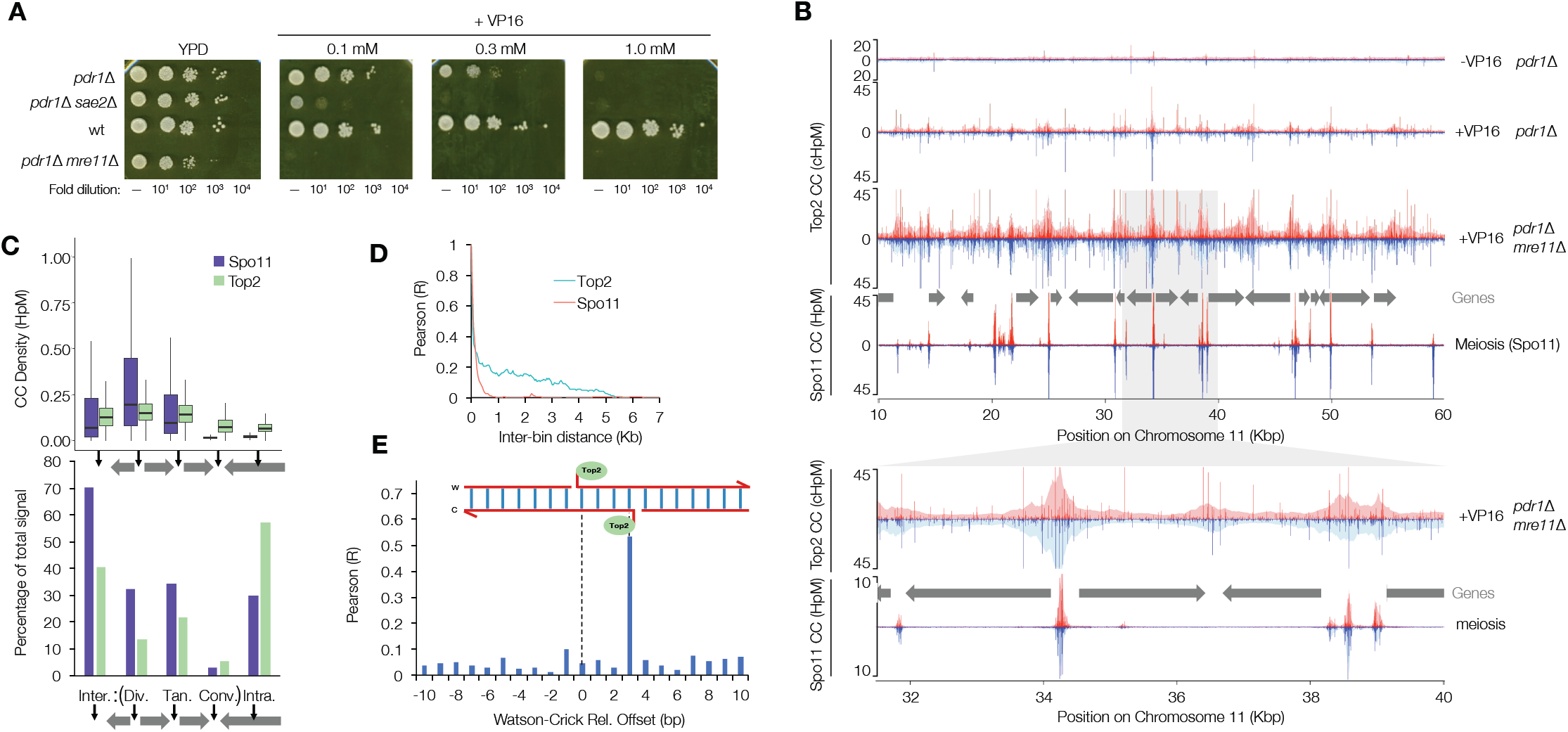
CC-seq maps covalent Top2-linked DNA breaks in *S. cerevisiae* cycling cells with nucleotide accuracy. **A)** Serial dilution spot tests of VP16 tolerance for the indicated strains. **B)** Nucleotide resolution mapping of *S. cerevisiae* Top2 CCs by CC-seq of the indicated strains after treatment for 4 hours with 1 mM VP16. Spo11-CC data is plotted for comparison. Top2 CC data were calibrated using a human DNA spike-in (**Methods**). Red and blue traces indicate CC-linked 5’ DNA termini on the Watson and Crick strands, respectively. Grey arrows indicate positions of gene open reading frames. Lower panels show an expanded view of the region from 31.5 to 40 Kbp. **C)** Quantification of Top2 and Spo11 CC signal stratified by genomic region. The genome was divided into intra and intergenic regions; the intergenic region was further divided into divergent, tandem and convergent based on orientation of flanking genes. Spo11 and Top2 activity mapped by CC-seq is expressed as box-and-whisker plots of density (upper and lower box limit: 3^rd^ and 1^st^ quartile; bar: median; upper and lower whisker: highest and lowest values within 1.5-fold of the interquartile range), or as the percentage of total mapped reads. **D)** Local correlation of Top2 or Spo11 CC-seq signals. Top2 or Spo11 CC-seq data were binned at 50 bp resolution and the Pearson correlation calculated between bins of increasing separation. **E)** Pearson correlation (r) of Top2 CC-seq signal on Watson and Crick strands, offset by the indicated distance. HpM = Hits per million mapped reads per base pair.

Next, we prepared sequencing libraries from mitotically growing etoposide-treated control, *sae2*Δ, and *mre11*Δ *S. cerevisiae pdr1Δ* cells (**Table S2**). A human DNA spike-in was included to enable calibration of relative signal between strains (**Methods**). Replicate libraries displayed high reproducibility (r > 0.89; **Figures S2A-C**), so were pooled. In untreated control cells, weak Top2 CC signal was distributed homogeneously across the genome, with few strong peaks (**Figure 2B**). Upon etoposide treatment, sharp single-nucleotide peaks arose at similar locations in all strains (**Figure 2B and S2D-E**), but with increased amplitude in the repair mutants—thereby directly linking the MRX pathway to repair of Top2 CCs in *S. cerevisiae*.

Global analyses revealed Top2 activity in *S. cerevisiae* to be relatively enriched in divergent and tandem intergenic regions (IGRs) and depleted in intragenic and convergent IGRs (**Figure 2B and C**), similar to, but less pronounced than, the patterns of Spo11 (**Figure 2B and C**). Such connections to global gene organisation are likely driven by the need for Spo11 and Top2 to interact with the DNA helix—an interpretation underpinned by an anticorrelation with nucleosome occupancy (**Figure S1D and S3A**). Nevertheless, the fact that *S. cerevisiae* Top2 CC signal is not correlated with proximal gene expression—neither in untreated cells, nor following etoposide exposure (**Figure S3B**)—suggests that in the gene-dense *S. cerevisiae* genome topological stress may be dealt with by Top2 at sites dislocated from where it is generated. By contrast, Spo11 activity is positively correlated with gene expression (**Figure S1E**), perhaps due to the influence transcription has upon the generation of higher order chromatin loop structures (Schalbetter et al., 2018), which in turn are known to pattern Spo11-DSBs (Sun et al., 2015).

Interestingly, Top2 CC signal remained weakly correlated over greater distances than Spo11 (**Figure 2D**), suggesting localised 5-10 kb wide regions of enriched activity—much larger than Spo11-DSB hotspots (generally <500 bp in size; **Figure 1C**; (Pan et al., 2011))— suggesting factors other than just nucleosome occupancy influence local Top2 catalysis. Top2 CCs are strongly correlated between Watson and Crick strands when offset by 3 bp (**Figure 2E and S2F-K**)—as expected for the 4 bp overhang generated by Top2-DSBs— further demonstrating the nucleotide resolution accuracy of CC-seq. Together these data describe how CC-seq reveals a strand-specific nucleotide resolution map of etoposide-induced Top2 CCs.

### CC-seq maps TOP2 CCs in the Human genome

Having demonstrated that CC-seq is applicable for mapping two divergent types of topoisomerase-like covalent complexes in *S. cerevisiae*, we applied CC-seq to map TOP2 activity in a human cell system. Asynchronous, sub-confluent RPE-1 cells were treated with etoposide (VP16) in the presence of MG132 proteasome inhibitor, in order to limit potential proteolytic degradation of TOP2 CCs that might otherwise hamper enrichment (**Figure 1A**). Slot-blotting and immunodetection with anti-TOP2β antibody demonstrated complete recovery of input TOP2β-linked CCs within the column elution, with no TOP2β remaining in the flow through or being removed by the washes (**Figure 3A**). Eluted material was used to generate sequencing libraries from three replicate control (–VP16) and four replicate etoposide-treated samples (+VP16), and high-depth paired-end sequencing reads (**Table S2**) were aligned to the human genome (hg19; **Methods**).

**Figure 3.**
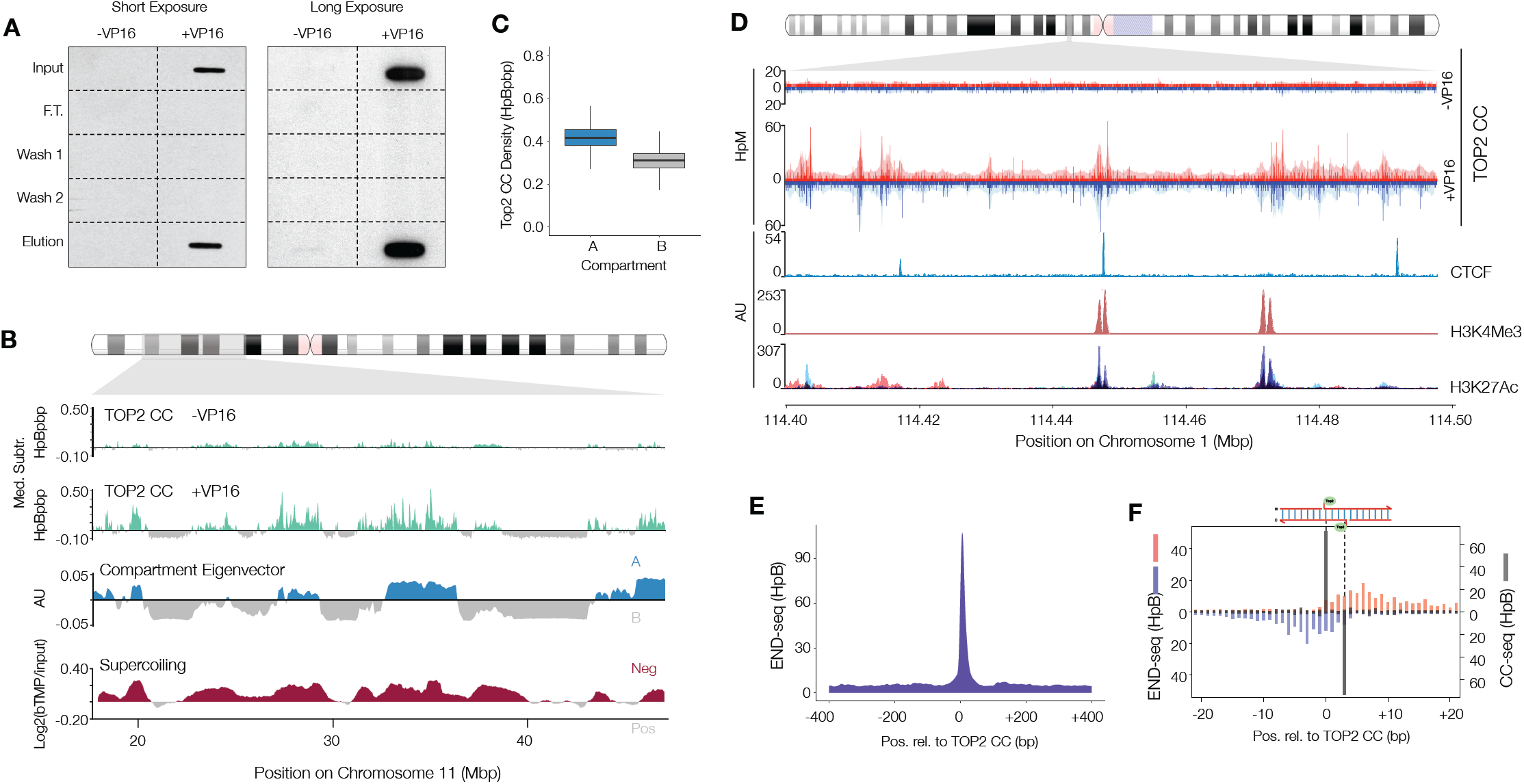
CC-seq maps TOP2-linked DNA breaks in Human cells with nucleotide accuracy. **A)** Anti-TOP2β western slot blot of input, flow through, wash, and elution fractions. RPE-1 cells were treated or not with VP16, prior to processing according to **Figure 1A**. **B)** Broad-scale maps of *H. sapiens* TOP2 CCs produced by CC-seq in RPE-1 cells ±VP16. Raw data were scaled, binned, smoothed and median subtracted prior to plotting (**Methods**). Chromatin compartments revealed by Hi-C eigenvector analysis (Darrow et al., 2016), and supercoiling revealed by bTMP ChIP-seq (Naughton et al., 2013) are shown for comparison. **C)** Quantification of TOP2 CC in chromatin compartments A and B. Data are expressed as box-and-whisker plots of density as for **Figure 2C**. **D)** Fine-scale mapping of *H. sapiens* TOP2 CCs by CC-seq in RPE-1 cells ±VP16. Red and blue traces indicate TOP2-linked 5’ DNA termini on the Watson and Crick strands, respectively. Pale shaded areas are the same data smoothed according to local density (see **Methods**). RPE-1 CTCF and H3K4Me3 ChIP-seq data plus H3K27Ac ChIP-seq data overlaid from seven cell lines (ENCODE, 2012) is shown for comparison. **E)** Medium-scale aggregate of END-seq mapped DSBs (Canela et al., 2017) surrounding nucleotide resolution CC-seq mapped TOP2 CCs. **F)** Fine-scale strand-specific aggregate of END-seq mapped DSB ends (red and blue bars) and nucleotide resolution CC-seq mapped TOP2 CCs (grey bars) surrounding strong TOP2 CC sites.

Replicates displayed high correlation in the distribution of TOP2 CC signal at broad scale (r values ≥ 0.79; **Figure S4A-B**), and so were pooled. Visual inspection revealed that -VP16 and +VP16 signals are spatially-correlated (**Figure 3B**), but that +VP16 signal intensity is less uniform, with more dynamic range (**Figure S4C-D**). These differences are likely due to higher signal-to-noise enabled by enrichment of TOP2 CC following VP16 treatment. Like yeast Top2 libraries (**Figure 2E**), nucleotide resolution analysis of human CC-seq libraries displayed a skew towards Watson-Crick read-pairs that are offset by 3 bp, as expected for the 4 bp 5’ overhang generated by TOP2 (**Figure S4E**). This skew is much greater in VP16-treated samples (**Figure S4E**), supporting the view that +VP16 libraries have greater Top2 CC enrichment.

The human genome encodes two type-IIA topoisomerases, TOP2α and TOP2β (**Introduction**). To demonstrate the specificity of CC-seq to map human TOP2 activity we used CRISPR-Cas9 to generate a human RPE-1-derived cell line with homozygous knockout mutations in the non-essential *TOP2B* gene, and arrested cells in G1 (**Figure S5A**) when TOP2α is not expressed (Heck et al., 1988; Woessner et al., 1991). TOP2α and TOP2β expression was undetectable in such G1-arrested cells (**Figure S5B**), and importantly, VP16-induced ?-H2AX foci were reduced ∼7-fold relative to wild type control lines, indicating that most of the signal is TOP2β-dependent (**Figure S5C-D**).

Next, we applied CC-seq to asynchronous and G1-arrested wild type and *TOP2B*^*-/-*^ cells. In G1, VP16 exposure induced localised regions of enriched TOP2 CC formation in wild type cells that were largely diminished or absent in *TOP2B*^*-/-*^ cells (**Figure S5E**). By comparison, TOP2β deletion did not prevent VP16-induced signal in asynchronous cells (**Figure S5E**), in which TOP2α is still present. Together, these results indicate that CC-seq detects a mixture of both TOP2α and TOP2β covalent complexes in wild type human cells depending on the cell cycle phase.

### Spatial distribution of Top2 activity in the human genome

The human genome is divided into a relatively gene-rich ‘A’ compartment and a relatively gene-poor ‘B’ compartment, the 3D spatial segregation of which can be determined by Hi-C (Lieberman-Aiden et al., 2009). In both untreated and VP16-treated asynchronous wild type cells, TOP2 CC are enriched in the active A compartment (**Figure 3B and C**), consistent with the role of TOP2 in facilitating transcription. TOP2 activity also correlated with regions of negative DNA supercoiling (Naughton et al., 2013) (**Figure 3B**). These observations provide a functional link between TOP2 activity, DNA topological stress, and large-scale chromatin compartments.

We next focused our attention in more detail on the genome-wide pattern of TOP2 activity enriched by VP16 treatment. Simple visual inspection (**Figure 3D**) suggested that TOP2 CC signals are enriched around genome features previously identified as regions of high TOP2-binding and etoposide-induced DNA strand breakage, including sites of CTCF-binding (Uusküla-Reimand et al., 2016; Canela et al., 2017) and active transcription start sites (TSSs) marked by H3K4Me3 and H3K27Ac (Baranello et al., 2014; Yang et al., 2015).

Genomic maps of etoposide-induced DNA breaks were recently generated using a related methodology, ‘END-seq’ (Canela et al., 2017), that is not specific for protein-linked TOP2 CC. Aggregating END-seq data around strong CC-seq positions revealed concordant peak signals at broad scale (**Figure 3E**), demonstrating a general agreement in the positions reported by the two techniques. However, at finer scale, END-seq positions are offset by 1-15 bp in the 3’ direction relative to CC-seq positions on each strand (**Figure 3F**)— something not observed when aggregating CC-seq data around the same loci (**Figure 3F**). This important difference is likely to be explained by the different populations of DNA breaks mapped by each technique: specific TOP2 cleavage positions (CC-seq) versus DNA breaks that have undergone limited nucleolytic resection (END-seq).

To investigate the relationship between CTCF and TOP2 activity, we filtered CTCF motifs to include only those that overlapped an RPE-1 CTCF ChIP-seq peak (**Methods;** (Akimitsu et al., 2003)). After aligning Watson and Crick motifs in the same orientation, we frequently observed a strong TOP2 CC peak upstream of the motif centre and accompanied by more distal, weaker peaks on both sides of the motif (**Figure 4A and S6A**). Loci containing multiple CTCF-binding motifs display more complex TOP2 CC patterns (**Figure 4A and S6B**) with enrichment on both sides of the double motif, as would be expected from the aggregation of a heterogeneous population of CTCF and TOP2 activity present across different cells.

**Figure 4.**
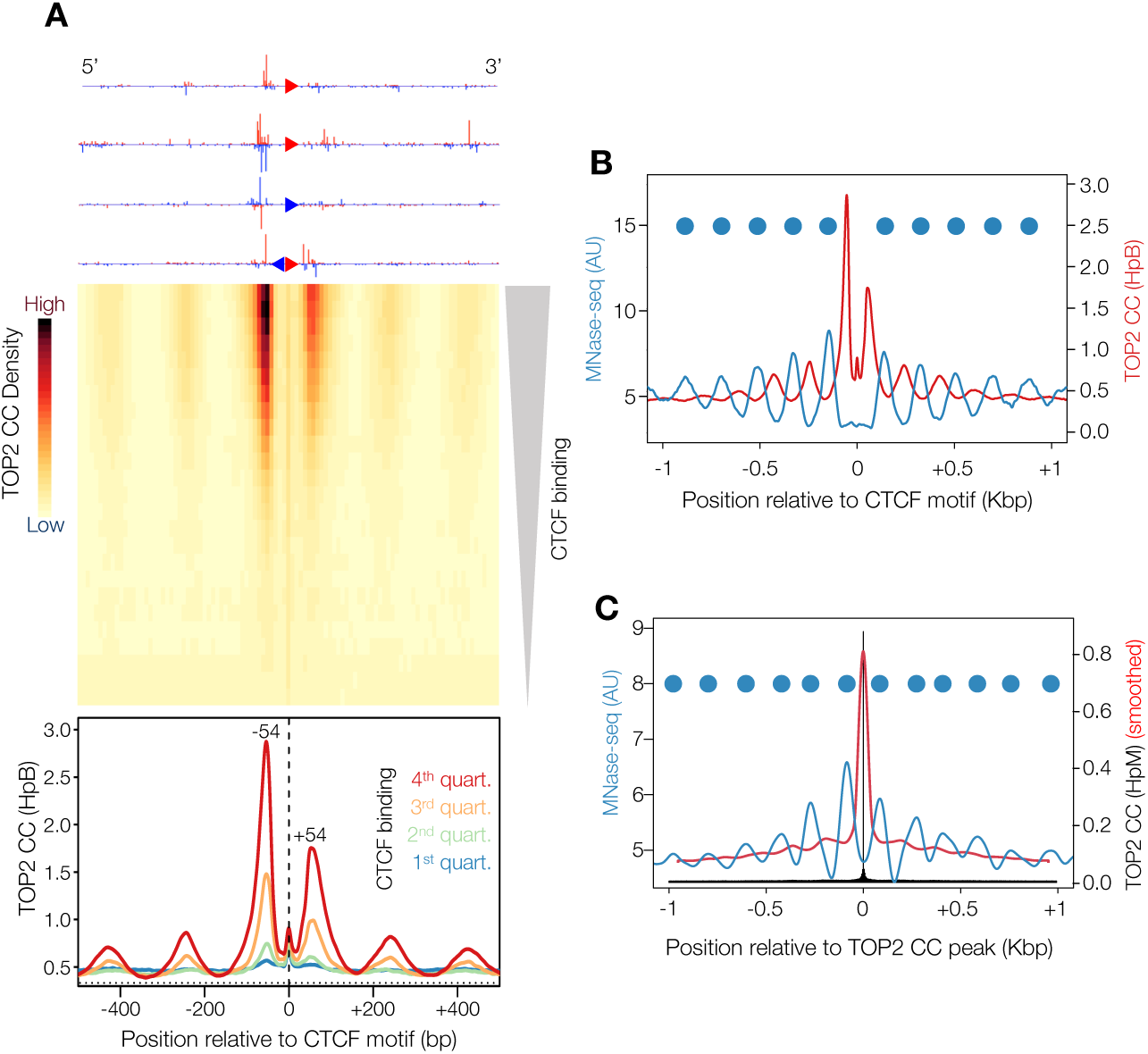
CTCF-proximal TOP2 activity is tightly confined within nucleosome-depleted regions. **A)** Aggregation of TOP2 CCs in a 1 Kbp window centred on orientated CTCF motifs in human RPE-1 cells. Four example CTCF loci are shown orientated in the 5’-3’ direction (top). A heatmap of all CTCF-motifs in the human genome, with 25 rows stratified by the strength of colocalising CTCF ChIP-seq peaks in RPE-1 cells (middle). The colour scale indicates average TOP2 CC density. Motifs are also stratified into 4 quartiles of CTCF-binding, and the average TOP2 CC distribution in each quartile plotted (bottom). **B)** Aggregated TOP2 CC distribution (red line) in the highest quartile of CTCF-binding compared with the average MNase-seq signal (blue line). **C)** Aggregated TOP2 CC distribution (black and red lines: single-nucleotide resolution and smoothed, respectively) and the average MNase-seq signal (blue line) surrounding strong TOP2 CC sites. In (B) and (C) peaks in MNase-seq signal indicate inferred nucleosome positions (blue circles).

We next stratified CTCF motifs into quantiles based on proximal CTCF binding intensity, and aggregated CC-seq signals centred on these loci (**Figure 4A**). Most Top2 CC signal is concentrated in two peaks located ±∼54 bp relative to the centre of the CTCF motif, with the upstream peak ∼2-fold more intense (**Fig 4A, middle and bottom panels**). Remaining Top2 CC signal is focused in a flanking array of weaker peaks. This signal pattern has been reported recently for VP16-induced DSBs (Canela et al., 2017), and has been suggested to result from a role for TOP2 in facilitating genome organisation by cohesin-dependent loop extrusion operating in the context of CTCF and other boundary elements. In support of this interpretation, total aggregated TOP2 CC signal across these regions is positively correlated with CTCF occupancy as measured by ChIP-seq (**Figure 4A**); interestingly however, TOP2 CCs were not correlated with interaction strength of associated loops (**Figure S6C**), annotated previously by Hi-C (Darrow et al., 2016). The positions of TOP2 CC surrounding CTCF motifs anticorrelate with the well-positioned nucleosomes at these sites (**Figure 4B**). However, such anticorrelation with nucleosome positioning is not a unique feature of CTCF loci, but is instead a general property of all TOP2 CC sites (**Figure 4C**), consistent with maps of both Top2 CC (**Figure S3B**) and Spo11 (**Figure S1E;** (Pan et al., 2011)) in *S. cerevisiae*.

In human cells, etoposide-induced TOP2 CCs are also enriched around gene TSSs (**Figure 5**), typically concentrated in broad peaks ∼1-2 Kbp downstream and ∼0.5-1 Kbp upstream of the TSS (**Figure 5A**; (Yang et al., 2015)). Similar to elsewhere in the genome, in the region immediately downstream of the TSS (0-400 bp), TOP2 CC signal anticorrelated with nucleosome occupancy (**Figure S6D**). Stratifying TSSs based on gene expression microarray data (Ganem et al., 2014) revealed a strong positive correlation (**Figure 5A**)— something not identified when prior etoposide-induced DSB maps were compared to nascent transcription as measured by GRO-Seq (Canela et al., 2017). Thus, while human TOP2 activity is correlated with total transcription of the gene, it is not proportional to local RNA polymerase activity when assayed at finer scale—suggesting a model where, as postulated for *S. cerevisiae* (**Figure S3A**), superhelical stress is resolved at sites dislocated from the process that generates them. Indeed, when analysed with an identical pipeline, both TSS-proximal TOP2 CCs (**Figure 5A**) and etoposide-induced DSBs (Canela et al., 2017) positively correlate with gene expression (**Figure S6E**), indicating that this correlation is not due to technical differences in mapping methodologies. Based on the twin-domain model of DNA supercoiling, topological stress is expected to increase not just with gene expression level, but also with increasing gene length (Liu and Wang, 1987). In support of this idea, we observe a strong positive correlation between human TOP2 CC enrichment and gene length independent from gene expression (**Figure S7A**). Interestingly, no such correlation with gene length is present in *S. cerevisiae* (**Figure S7B**), further supporting a model whereby, in this compact gene-dense genome, Top2 activity in the IGR is neither directly linked to expression nor length of the proximal gene.

**Figure 5.**
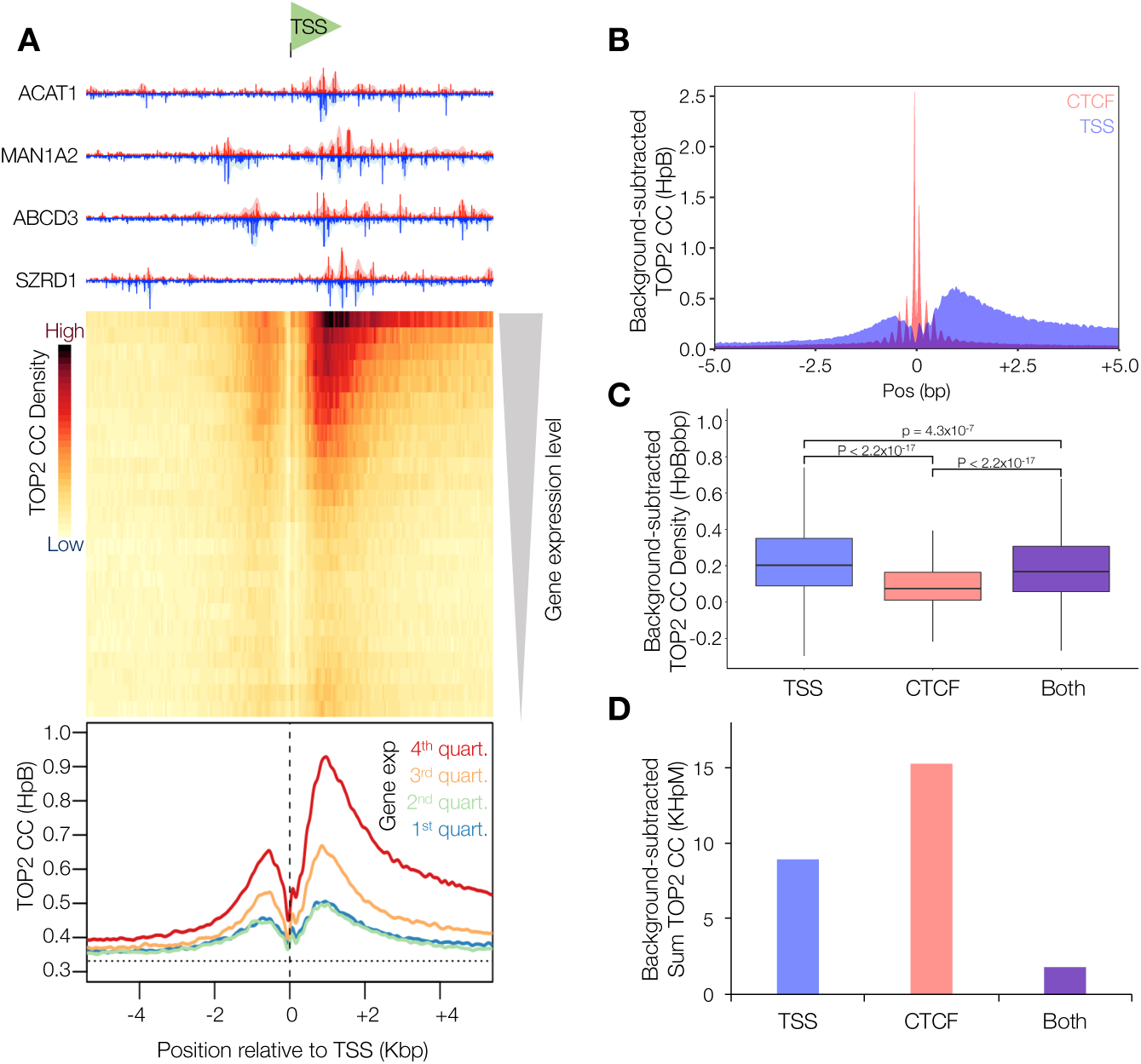
TSS-proximal TOP2 activity is strongly correlated with gene transcription. **A)** Aggregation of TOP2 CCs in a 10 Kbp window centred on orientated TSSs in human RPE-1 cells. Four example TSSs are shown orientated in the 5’-3’ direction (top). A heatmap of all TSSs in the human genome, with 25 rows stratified by gene expression level in RPE-1 cells (middle). The colour scale indicates average TOP2 CC density. Motifs are also stratified into 4 quartiles of gene expression, and the average TOP2 CC distribution in each quartile plotted (bottom). **B)** Comparison of CTCF-proximal and TSS-proximal TOP2 CC distributions in the highest quartile of CTCF-binding (pink) and gene-expression (blue), respectively. **C)** Average TOP2 CC density in 10 Kbp regions centred on the highest quartile of TSS, CTCF, or regions where both features are present. Data are expressed as box-and-whisker plots of density as in **Figure 2C**. Statistical significance was determined using the KS test. **D)** As in (C) but sum total TOP2 CCs found in these regions.

Notably, the increased density of etoposide-induced human TOP2 CCs at strongly expressed genes is 2.7-fold greater than at strongly bound CTCF motifs (**Figure 5B and C**). Thus, whilst strongly-bound CTCF motifs (14,924) greatly outnumber active genes (4,194)—leading, globally, to more TOP2 activity in total at CTCF sites (**Figure 5D**)—our data reinforce the important role of topoisomerase activity during transcription.

### Local DNA sequence and methylation status direct the formation of Top2 CCs

Previous reports suggest that etoposide induces a majority of SSBs, rather than DSBs, due to independent poisoning of each active site in the Top2 homodimer (Bromberg et al., 2003). In our yeast and human datasets, we similarly observe more single-strand Top2 CCs (90.1% and 94.2%, respectively) than double-strand Top2 CCs (9.9% and 5.8%, respectively) following etoposide exposure (**Figure 6A and S3C**).

**Figure 6.**
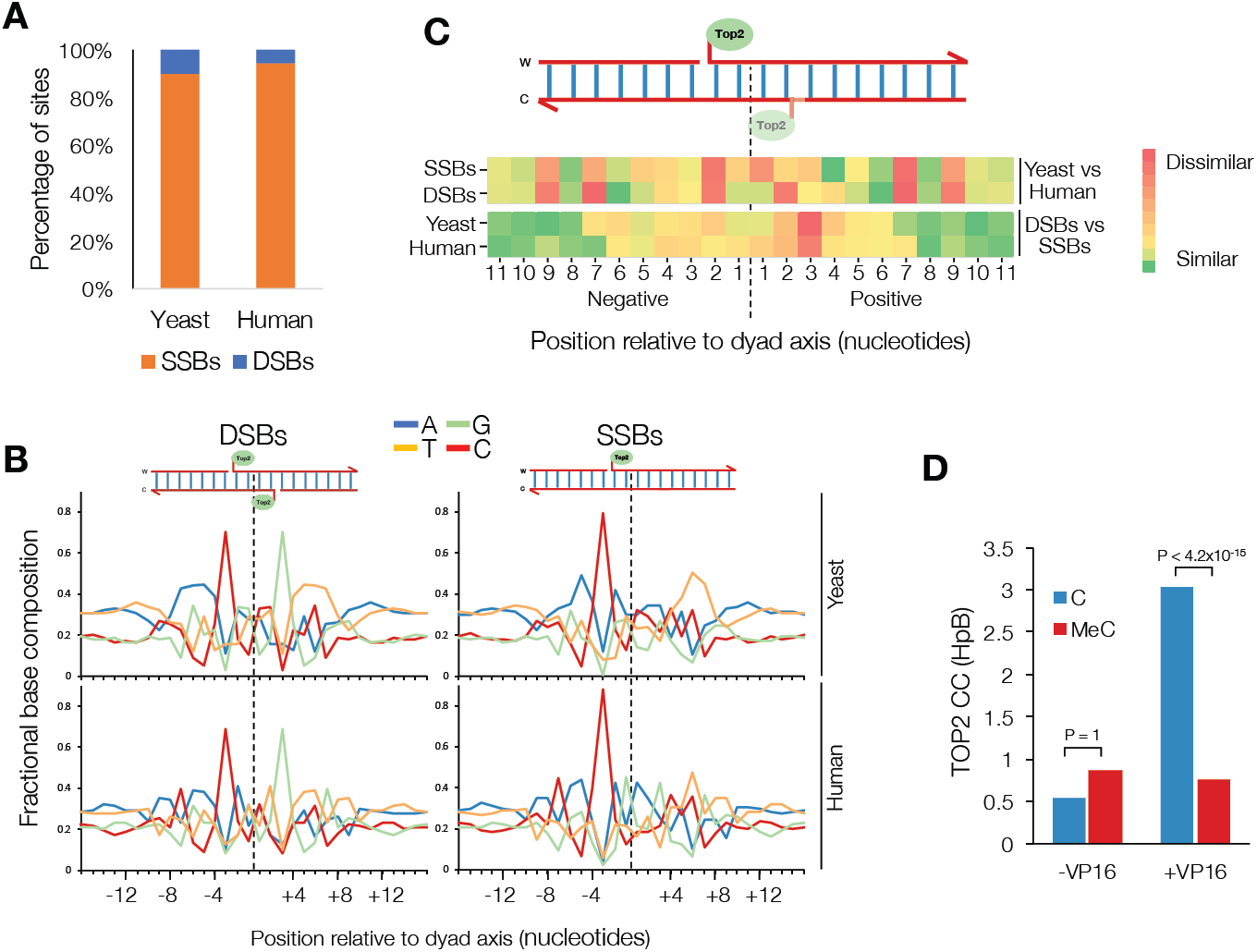
Local DNA sequence and methylation status direct the formation of Top2 CCs. **A)** The percentage of strong sites that are SSBs and DSBs in etoposide-treated *S. cerevisiae* and human cells. **B)** Average nucleotide composition over a 30 bp window centred on the DSB or (inferred) SSB dyad axis, in etoposide-treated *S. cerevisiae* and human cells. Values reported are for the top strand only. **C)** Heatmaps describing pairwise similarity of nucleotide composition patterns shown in (B). Rows 1-4 are the absolute differences between: yeast and human SSB patterns, yeast and human DSB patterns, yeast SSB and DSB patterns, and human SSB and DSB patterns, respectively. **D)** The average number of TOP2 CCs at the +1 position relative to methylated and unmethylated cytosines in human RPE-1 cells, ±VP16. Statistical significance was determined using KS test.

Prior, fine-scale mapping of Top2 cleavage at specific substrates indicated that cleavage patterns are influenced by local DNA sequence composition (Pommier et al., 1991; Strumberg et al., 1999). To investigate the generality of such findings, and to precisely determine how DNA sequence influences Top2 site preference across the yeast and human genome, we computed the nucleotide resolution average base composition around Top2 CC SSB and DSB sites (**Fig 6B**)—something made possible by the positional accuracy of our mapping technique. For DSBs, average DNA sequence patterns are rotationally symmetrical around the dyad axis of cleavage (**Figure 6B, left)**, consistent with the known homodimeric nature of eukaryotic Top2 (Pommier et al., 2016). Base positions with notable skews are restricted to the region ±11 bp from the dyad axis (p<10^−4^, Chi-squared test), consistent with the DNA contacts made by Top2 (Lee et al., 1989; Thomsen et al., 1990; Wu et al., 2011), yet with minor, species-specific differences at a few positions (±2, ±7, ±9; **Figure 6C, top**), suggesting subtle differences in the ScTop2 and HsTOP2 DNA binding surface. Positions −3 and +3 are the bases immediately 5’ to the scissile phosphodiester bonds cleaved by Top2 (**Figure 6B**). In the averaged DSB sequence motif, strong preference for cytosine at these positions (69.0% of DSBs) on the scissile strand (**Figure 6B, left)** agrees with *in vitro* studies of Top2 mechanism—attributed to favourable base-stacking interactions of etoposide with guanosine at the −1 position on the non-scissile strand (Pommier et al., 1991; Strumberg et al., 1999; Wu et al., 2011).

Importantly, whilst SSBs display similar overall symmetrical sequence features as DSBs, SSBs differed substantially at position +3, where the strong preference for guanine on the scissile strand (and paucity of cytosine on the non-scissile strand) is absent (**Figure 6B, right, and Figure 6C, bottom**). Furthermore, the preference for cytosine at the −3 position on the scissile strand increases to 88.1% for SSBs. This clear difference in sequence bias provides strong evidence that such positions truly reflect sites of preferred VP16-induced SSBs vs DSBs *in vivo*.

The human genome is subject to methylation of cytosine (meC) at CpG sites. Dramatically, meC completely abolishes the formation of VP16-induced TOP2 CCs (**Figure 6D**), suggesting this bulky group may interfere sterically with the VP16-Top2-DNA ternary complex, modifying TOP2 catalysis. Collectively, these observations demonstrate that local DNA sequence composition—including epigenetic marks—are major determinants of both TOP2 CC abundance and the differential generation of SSBs and DSBs by VP16 *in vivo*.

## Discussion

Topoisomerases are ubiquitous gatekeepers to the genetic material—facilitating changes to DNA topology, nuclear organisation, and chromosome segregation that are essential for the generation and propagation of all life on Earth. Yet, topoisomerase dysfunction can also prove harmful, generating difficult to repair protein-linked DNA breaks that create risks to genome stability. Thus, understanding how, where, and when topoisomerase activity takes place is of great importance. Here, we demonstrate a robust, technically simple and sensitive method to map sites of preferred type IIA topoisomerase activity across the eukaryotic genomes of yeast and human cells at strand-specific nucleotide resolution. Our methodology joins an extensive toolkit of complementary NGS solutions (Canela et al., 2017; Crosetto et al., 2013; Yan et al., 2017), each with its own set of unique advantages. However, many of these techniques do not achieve nucleotide resolution, either due to the use of nucleolytic processing to blunt 5’ DNA termini (Canela et al., 2017), and/or inability to directly map DNA breaks covalently linked to protein (Canela et al., 2017; Crosetto et al., 2013; Yan et al., 2017). Here, by utilising the tyrosyl phosphodiesterase activity of recombinant Tdp2 to unblock 5’-DNA termini non-nucleolytically, we are able to accurately map the position of not just Top2 CCs, but also related classes of topoisomerase-like enzymes such as Spo11, which acts uniquely in meiotic cells. Importantly, such specificity in our cleavage maps enable us to corroborate and extend—with completely independent methodology—prior research findings from meiosis (Pan et al., 2011), the impact that VP16 has on SSB versus DSB formation—and for the first time demonstrate the influence that primary DNA sequence composition and methylation status has on the process of Top2 catalysis *in vivo* in both yeast and human cells.

Top2 activity has long been associated with transcriptional activity (Yang et al., 2015; Brill et al., 1987; Baranello et al., 2014; Yu et al., 2017). In addition, recent research has identified etoposide-induced DSBs that colocalise with a subpopulation of chromatin-bound TOP2 on the loop-external side of CTCF-and Rad21-bound CTCF motifs (Canela et al., 2017; Uusküla-Reimand et al., 2016). TOP2 activity at these sites has been proposed to facilitate genome organisation driven by chromatin loop extrusion, and thereby to constitute a major fraction of the total nuclear TOP2 activity (Canela et al., 2017). Moreover, contrary to expectations, the frequency of etoposide-induced DSBs was found to be locally uncorrelated with nascent RNA levels (Canela et al., 2017)—generating some confusion in the understanding of where TOP2 activity is most prevalent and most important. Here, our independent CC-seq method and analyses permit us to revisit this question in explicit detail, and clarify the role of TOP2 in these processes. Importantly, whilst Top2 activity is certainly highly enriched at CTCF-bound genomic loci, the total amount of activity at any given CTCF locus is many times lower than observed at transcriptionally active genes. Nevertheless, it is critical to emphasise that because of the great abundance of CTCF-bound sites compared to active genes, total aggregated TOP2 activity is globally similar for CTCF-associated versus transcriptionally active regions of the genome.

Our data also permit us to investigate how other genomic features influence Top2 activity. For example, at both TSS and CTCF-proximal regions—and indeed in the wider genome of both yeast and human—Top2 cleavages are highly enriched in nucleosome-free DNA, in support of prior findings made at a subset of genomic loci (Capranico et al., 1990; Udvardy and Schedl, 1991; Käs and Laemmli, 1992). Notably, whilst yeast Top2 activity is also enriched near TSSs, it is concentrated immediately (120-140 bp) upstream, unlike human TOP2 activity, which is primarily concentrated in a broader peak 1-2 kb downstream, within the gene body. Furthermore, unlike human TOP2, yeast Top2 activity is uncorrelated with downstream gene expression. These differences may stem from the very different scale and transcriptional organisation of the yeast and human genomes, and the strong patterning of Top2 activity driven by accessibility to nucleosome-free intergenic promoter regions in *S. cerevisiae*. In support of this interpretation, similar nucleosomal constraints appear to substantially govern the activity of the related topoisomerase-like enzyme, Spo11, during meiosis (Pan et al., 2011) (**Figure S1** herein).

Despite these differences in distribution of Top2 activity at this larger scale, the DNA sequence preferences for yeast and human Top2 are similar, implying substantial conservation of the DNA binding interface, which is supported by analysis of the published crystal structure of VP16-TOP2-DNA (Wu et al., 2011). A nucleotide resolution genome-wide map of *E. coli* gyrase activity was recently reported (Sutormin et al., 2018)), revealing a sequence motif composed of a central 35 bp region containing the cleavage site, plus two flanking 47 bp regions with ∼10.5 bp periodicity centred ∼±40 bp from the site of cleavage, which are attributed to the known wrapping of proximal DNA around the gyrase C-terminal domains (Reece and Maxwell, 1991). Strikingly, whilst we observe a core central motif for Top2, there are no such flanking motifs over these ranges, in agreement with its lack of DNA wrapping domains (**Figure S3D**). We do, however, observe two more-proximal flanking regions centred ∼±20 bp from the site of cleavage, each with ∼10.5 bp periodicity (**Figure S3E**), suggesting that Top2 also bends its DNA substrate—in agreement with a recent biophysical study (Huang et al., 2017).

We have used CC-seq to probe MRX-dependent repair of Top2 CCs in *S. cerevisiae*, corroborating and extending prior genetic experiments in this organism (Hamilton and Maizels, 2010; Stepanov et al., 2008), and others (Hoa et al., 2016; Hartsuiker et al., 2009). Importantly, calibration of our signal using an internal human standard enables the relative quantification of Top2 CCs, and thus directly demonstrates for the first time that loss of Mre11 activity increases cellular levels of Top2 CCs in *S. cerevisiae*. The antigen-independent enrichment step allows calibration to be performed without genetic epitope-tagging or reconstitution of recombinant protein-DNA complexes, making CC-seq ideally suited to the future study of repair dynamics.

We have demonstrated that CC-seq maps CCs of both human TOP2α and TOP2β— proteins that are conserved amongst all vertebrates and whose functional differences remain a subject of interest within the field (Austin et al., 2018). Given the essential requirement for TOP2α during mitosis (Akimitsu et al., 2003), and the apparent inability of TOP2β to compensate for its loss (Grue et al., 1998), it will be particularly interesting to determine the genomic distribution of TOP2α/TOP2β activities throughout the cell cycle, and to relate this to 3D chromatin conformation measured by Hi-C. Whilst not essential for cell proliferation, TOP2β plays an important role in promoting transcriptional programmes associated with neuronal development (Tiwari et al., 2012), and this function cannot be supported by TOP2α. Furthermore, loss of TDP2 function inhibits TOP2-dependent gene transcription and leads to neurological symptoms including intellectual disability, seizures and ataxia (Gómez-Herreros et al., 2014). Thus, accurate genomic maps of TOP2β CC formation in these tissues will help to identify those genes most pertinent to the development of neurodegeneration. Moreover, as demonstrated by our Spo11 maps, the antigen-independent, high-efficiency protein-DNA enrichment process makes CC-seq generally applicable for mapping not just Top2 activity in diverse organisms, but also for mapping similar types of topoisomerase-DNA covalent complexes, or even those less distinct DNA-protein crosslinks that arise upon ablation of the DNA-dependent protease SPRTN/Wss1 (Stingele et al., 2015).

## Acknowledgements

We thank Keith Caldecott, Antony Oliver and Peter Hornyak for sharing recombinant TDP2, and Jon Nitiss for sharing the *pdr1*Δ strains.

## Author Contributions

W.G., D.J. and M.J.N. conceived the project and developed the CC-seq methodology. W.G., D.J., H.T. and R.A. generated whole genome CC-seq libraries. W.G. performed all data processing and analysis, and all human cell work. D.J., H.T. and R.A. performed yeast sensitivity and meiotic DSB assays. T.J.C. developed the mapping pipeline and associated tools. W.G. and M.J.N. interpreted the observations and wrote the manuscript.

## Data availability

All strains and cell lines listed in Table S3 and S4 are available on request. Sequencing reads are available via the NCBI Sequence Read Archive (accession numbers pending).

## Methods

### Yeast strains, culture methods and treatment

The *Saccharomyces cerevisiae* yeast strains used in this study are described in **Table 2** and were derived using standard genetic techniques. Strains used for Spo11 mapping are isogenic to the SK1 subtype and carry the *sae2*Δ∷kanMX gene disruption allele. Strains used for Top2 mapping are isogenic to the BY4741 subtype and carry the *pdr1DBD-CYC8* drug sensitivity cassette (Stepanov et al., 2008). For Spo11 DSB mapping, cells were induced to undergo synchronous meiosis as follows: Cells were grown overnight to saturation in 4 ml YPD medium (1% yeast extract, 2% peptone, 2% glucose supplemented with 0.5 mM adenine and 0.4 mM uracil) at 30 °C, then diluted to OD_600_ 0.2 in 200 ml YPA medium (1% yeast extract, 2% peptone, 1% potassium acetate) and grown vigorously for 15 h at 30 °C. Cells were then washed with water, resuspended in 200 ml sporulation medium (2% potassium acetate supplemented with diluted amino acids), incubated vigorously for 6 h at 30 °C and harvested by centrifugation. For etoposide treatment, cells were grown overnight to saturation in 4 ml YPD medium at 30 °C, then diluted to OD_600_ 0.5 in 100 ml YPD and grown until OD_600_ 2. 50 ml cultures were then incubated for a further 4 h in the presence of either 1 mM etoposide or 2% DMSO, before harvesting by centrifugation.

### Human cell lines, culture methods and treatment

The human cell lines used in this study are described (**Supplementary Table 3**). Human hTERT RPE-1 cells were obtained from ATCC and cultured at 37 °C, 5% CO2 and 3% O2 in Dulbecco’s Modified Eagle’s Medium DMEM/F-12 (Sigma), supplemented with 10% Fetal Calf Serum (FCS) and 90 Units/ml Penicillin-Streptomycin. For all experiments (CC-seq, Slot Blot, WB, IF, FACs) with asynchronous cell populations, RPE-1 cells were seeded at a density of (3.5×10^3^ cells/cm^2^l) and incubated for 72 h at 37 °C, to ensure subconfluent log-phase growth (∼70% confluency) at the time of the experiment. For G1 cell populations, WT and *TOP2B*^*-/-*^ RPE-1 cells were seeded at a density of 5×10^3^ cells/cm^2^ and incubated for 48 h at 37 °C in DMEM/F12 containing 10% FCS, prior to a further 24 h incubation in DMEM/F12 medium containing no FCS to ensure complete G1-phase arrest (verified by FACs, see below). For experiments with proteasome inhibitor and etoposide (CC-seq, Slot Blot), cells were preincubated with 5 μM MG132 (Sigma) for 90 min, trypsinised and incubated in suspension with 5 μM MG132 and 100 μM etoposide (Sigma) for 20 min at 37 °C. For experiments with etoposide alone (IF), adherent cells were treated with 100 μM etoposide for 20 min at 37 °C.

### Generation of *TOP2B-/-* RPE-1 cells

The oligonucleotides 5’-CACCGCCGCAGCCACCCGACT and 5’-AAACAGTCGGGTGGCTGCGGC (identified using Benchling; https://benchling.com) were annealed and cloned into pX330 following BbsI restriction, as described previously (Hsu et al., 2013). This SpCas9/ trugRNA co-expression plasmid was transiently expressed in RPE-1 cells to target the 17 bp target sequence GCCGCAGCCACCCGACT (TGG) within exon 1 of *TOP2B*. Single clones were trypsinised and passaged to isolated culture vessels prior to screening for absent protein by Western Blot (WB). All experiments involving *TOP2B*^*-/-*^ were conducted using the RPE-1 clone T2B/6, in which no TOP2β is detectable by WB or IF.

### Spot tests of chronic etoposide sensitivity

Single colonies of each *S. cerevisiae* strain were incubated overnight in 4 ml YPD at 30 °C with shaking. 0.1 ml of this starter was used to inoculate 4 ml YPD, prior to incubation for 5 hours at 30 °C. Cultures were diluted to make a stock with an OD_600_ of 2.0, then this was 10-fold serially diluted five times. Each dilution was spotted onto plates containing 0, 0.1, 0.3 or 1.0 mM VP16, prior to incubation for three days at 30 °C. Plates were imaged on a 2400 Photo scanner (Epson).

### Southern blotting of meiotic Spo11 DSBs

Approximately 2 μg of genomic DNA (isolated by non-proteolysing Phenol-Chloroform extraction, as described below) was digested at 37 °C overnight using *Pst*I restriction enzyme (NEB) in NEBuffer 3.1 (100 mM NaCl, 50 mM Tris Base.HCl pH 7.9, 10 mM MgCl2, 100 μg ml-1 BSA). Additional *Pst*I was added for 4 hours before the addition of NEB purple loading dye to 1×. Digested samples were proteolysed using 1 mg ml-1 Proteinase K (Sigma) at 60 °C for 30 minutes, left to reach room temperature before 10 μg was loaded on a 0.7% 1× TAE agarose gel (40 mM Tris Base.HCl, 20 mM glacial acetic acid, 1 mM EDTA pH 8.0) containing 50 μg ml-1 ethidium bromide. DNA was separated in 1× TAE at 60 V for 18 hours. The gel was imaged using InGenius (Syngene) bioimaging system to check migration and then exposed to 180 mJ/m^2^ UV in the Stratalinker (Stratagene). The gel was then soaked in three times its volume of denaturation solution (0.5 M NaOH, 1.5 M NaCl) for 30 minutes and then transferred to Zetaprobe (Bio-Rad) membrane by means of a vacuum at 55 mBar for 2 hours. After transfer the membrane was washed in water ten times and then cross-linked by exposing the membrane to 120 mJ/m^2^ UV in the Stratalinker. The membrane was incubated in 30 ml of hybridisation buffer (0.5 M NaHPO_4_ buffer pH 7.5, 7% SDS, 1 mM EDTA, 1% BSA) at 65 °C for 1 hour. The *MXR2* probe for looking at the *HIS4∷LEU2* locus was created from 50 ng of template DNA, 0.1 ng of Lambda DNA (NEB) digested with *BstE*II (NEB), and water. The mix was denatured at 100 °C for 5 minutes then put on ice. High Prime (Roche) was added in addition to 0.5-3 mBq of α-^32^P dCTP and incubated at 37 °C for 15 minutes. 30 μl 1× TE was added and the probe spun through a G-50 spin column (GE Healthcare) at 400 × *g* for 2 minutes. The probe was then denatured by incubating at 100 °C for 5 minutes and then put on ice before being added to 20 ml hybridisation mixture. The original 30 ml hybridisation buffer was discarded and the 20 ml containing the probe was added to the membrane and incubated overnight at 65 °C. After incubation, the membrane was washed five times with 100 ml pre-warmed Southern wash buffer (1% SDS, 40 mM NaHPO_4_ buffer pH 7.5, 1 mM EDTA) and exposed to phosphor screen overnight.

### Western blotting

Whole human cell extracts (WCE) were harvested by direct lysis in 1x Laemmli loading buffer, denatured for 10 min at 95 °C and sonicated for 30 s using Bioruptor^®^ Pico. Samples were subjected to SDS-PAGE (7% or gradient gel) and transferred to nitrocellulose membrane. Primary immunodetection was with antibodies targeting TOP2β (Clone 40, BD Biosciences), TOP2α (ab52934, Abcam), or Ku80 (ab80592, Abcam). Secondary immunodetection was with HRP-conjugated Rabbit anti-Mouse IgG (ThermoFisher), prior to detection of peroxidase activity using ECL reagent and X-Ray film (Scientific Laboratory Supplies Ltd).

### Slot blotting

Samples were diluted 4-fold (500 uL total volume) in NaPO4 buffer (25 mM, pH 6.5), and slot blotted onto 0.2 μM nitrocellulose membrane (Amersham), using the Minifold I (Whatman) manifold. The wells were washed twice with 750 μL NaPO4 buffer. The membrane was then blocked with 10% milk-TBST for 1 h at RT, prior to incubation overnight with anti-TOP2β antibody (Clone 40, BD Biosciences) at 4 °C. The membrane was washed 4 times with TBST, incubated with HRP-conjugated Rabbit anti-Mouse IgG (ThermoFisher) for 1 h at RT, washed 4 times with TBST, and incubated with ECL detection reagent for 1 min. X-Ray film was used for detection.

### Fluorescence-Assisted Cell Sorting (FACS)

Approximately 10 million RPE-1 cells were trypsinised, washed once in PBS and resuspended in 1.5 ml PBS. 3.5 ml ethanol was added dropwise to pellet, with vortexing. Cells were fixed for 1 hr at 4 °C, prior to centrifugation and aspiration of the supernatant. Cells were washed twice with PBS, prior to resuspension in 0.5 ml 0.25% Triton-X100-PBS for 15 min on ice. Cells were pelleted by centrifugation, supernatant was aspirated, and the pellet was resuspended in 0.5 ml TBS containing 10 ug/ml RNase A (Sigma) and 167 nM Sytox Green (ThermoFisher). After 30 min incubation in the dark at RT, the suspension was filtered through fine mesh into test tubes. DNA content in 50,000 cells was analysed using the Accuri C6 (BD Biosciences), with gating to exclude doublets and cell debris.

### Immunofluorescence

Cells were seeded onto glass coverslips (Agar Scientific), incubated and treated according to the protocol outlined above. Cells were then fixed with 4% paraformaldehyde PBS for 10 min, washed three times with PBS, permeabilised with 0.2% Triton-X100-PBS for 10 min, blocked for 1 h with 10% FCS-PBS, incubated with primary antibodies targeting Phospho-Histone H2AX (S139) (JBW-301, Merck Millipore) and TOP2α (ab52934, Abcam), washed three times with PBS, incubated with secondary antibodies Alexa 488-conjugated Goat anti-mouse IgG (Fisher) and Alexa 647-conjugated Goat anti-Rabbit (Fisher), washed three times with PBS, washed once with distilled water, and mounted with VECTASHIELD containing DAPI (Vector Laboratories).

### High-content microscopy

Automated wide-field microscopy was performed on an Olympus ScanR system (motorised IX83 microscope) with ScanR Image Acquisition and Analysis Software, 40x/0.6 (LUCPLFLN 40x PH) dry objectives and Hamamatsu ORCA-R2 digital CCD camera c4300. Numbers of anti-phospho-Histone H2AX (S139) foci (Alexa 488; FITC filter) were quantified in the nuclear region colocalising with DAPI, using Olympus ScanR Analysis software. TOP2α signal (Alexa 647; Cy5 filter) was also quantified in this region.

### CC-seq: Enrichment of protein-linked DNA

1.5×10^7^ RPE-1 cells were treated as described above, pelleted by centrifugation, washed once with 15 ml ice-cold PBS, and resuspended in three aliquots of 400 μL ice-cold PBS. 1×10^9^ yeast cells were treated as described above, then spheroplasted in 1.5 ml spheroplasting buffer (1 M sorbitol, 50 mM NaHPO_4_ buffer pH 7.2, 10 mM EDTA) containing 200 μg/ml Zymolyase 100T (AMS Biotech) and 1% β-mercaptoethanol (Sigma) for 20 min at 37 °C. 2 μL Protease Inhibitor Cocktail and 2 μL Pefabloc (Sigma) were added before splitting into five aliquots of 400 μL. All subsequent steps of the protocol are the same for yeast and human samples. 1 ml ice-cold ethanol was added to 400 μL cell suspensions in microcentrifuge tubes, mixed, incubated for 10 min on ice, and pelleted by centrifugation. The supernatant was thoroughly removed by aspiration, prior to addition of 200 μL 1x STE buffer (2% SDS, 0.5M Tris pH 8.1, 10 mM EDTA, 0.05% bromophenol blue), cell disruption using a pestle (VWR), addition of a further 400 μL 1x STE buffer, and incubation for 10 min at 65 °C. Samples were cooled on ice and 500 μL Phenol-Chloroform-isoamyl alcohol (25:24:1; Sigma) was added. The mixtures were emulsified by shaking and pipetting 5 times with a 1 ml micropipette, prior to phase separation by centrifugation at 20,000 g for 20 min. By minimising mechanical shearing of the lysate prior to phenol chloroform extraction, peptides that are covalently linked to high molecular weight DNA segregate into the aqueous phase. 500 μL of the aqueous phase was removed to a clean microcentrifuge tube and nucleic acids were precipitated with 1 ml ice-cold ethanol, pelleted by centrifugation, washed with ice-cold 70% ethanol, and dissolved in TE buffer overnight at 4 °C. Samples were then incubated with 0.2 mg/ml RNase A (Sigma) for 1 h at 37 °C; nucleic acids were precipitated with 1 ml ethanol, pelleted by centrifugation, washed twice with 70% ethanol and dissolved in TE overnight. Aliquots were combined to a total of 1 ml and sonicated to an average fragment size of 300-400 bp with Covaris (duty cycle: 10%, intensity/peak power incidence: 75W, cycles/burst: 200, time: 15 min). 1 ml of sonicated sample was added to 1.2 ml of binding buffer (10 mM Tris pH 8.1, 10 mM EDTA, 0.66 M NaCl, 0.22% SDS, 0.44% N-Lauroylsarcosine sodium salt). Each sample was divided over several Miniprep (QIAGEN) silica-fibre membrane spin columns, such that the total DNA loaded to each was approximately 20 μg. The flowthrough was reapplied to the column to improve binding. Columns were washed 6 times with 600 μL of TEN (10 mM Tris, 1 mM EDTA, 0.3 M NaCl) per 1 min wash, prior to elution with 100 μL TES (10 mM Tris, 1 mM EDTA, 0.5% SDS).

### CC-seq: DNA end repair and adapter ligation

Eluted products were pooled to 500 μL in TES and incubated with 1 mg/ml Proteinase K (Sigma) for 30 min at 60 °C, prior to overnight ethanol precipitation at −80 °C with 1.41 ml ethanol, 0.2 mg/ml glycogen and 200 mM NaOAc. The DNA-glycogen precipitate was pelleted by centrifugation at 20,000 g for 1 hr at 4 °C, washed once with 1.5 ml 70% ethanol, and re-pelleted by centrifugation. The supernatant was aspirated and the pellet was air-dried for 10 min at RT, prior to solubilisation in 52 μL 10 mM Tris-HCl. DNA concentration was measured in a 2 μL sample with the Qubit (ThermoFisher) and High Sensitivity reagents. The remaining 50 μL was used as input for one round of end repair and adapter ligation with NEBNext Ultra II DNA Library Preparation kit (NEB), according to manufacturer’s instructions, except for the use of a custom P7 adapter (see table). The use of custom adapters is to allow differentiation of the sheared end (P7 adapter) from the Top2/Spo11 end (P5 adapter). After ligation of the P7 adapter, DNA was isolated with AMPure XP beads (Beckman Coulter) according to manufacturer’s instructions (beads:input of 78:90) and eluted in 50 μL 10 mM Tris-HCl. Samples were diluted 2-fold with 50 μL Tdp2 reaction buffer (100 mM TrisOAc, 100 mM NaOAc, 2 mM MgOAc, 2 mM DTT, 200 μg/ml BSA) and incubated with 3 μL of 10 μM recombinant human TDP2 (Hornyak et al., 2016; Johnson et al., 2019), for 1 h at 37 °C. DNA was isolated again with AMPure XP beads (beads:input of 103:103) and eluted in 52 μL 10 mM Tris-HCl. Next, a second round of end repair and adapter ligation was conducted, using adapter P5 (see table). After ligation of the P5 adapter to the Top2/Spo11-cleaved end, DNA was isolated with AMPure XP beads (beads:input of 78:90) and eluted in 17 μL 10 mM Tris-HCl.

### CC-seq: PCR and size selection

DNA concentration was measured in 2 μL using the Qubit. The remaining 15 μL was used as template for the PCR step of the NEBNext Ultra II PCR step using universal primer (P5 end) and indexed primers for multiplexing (P7 end), according to manufacturer’s instructions. PCR reactions were diluted with 50 μL 10 mM Tris-HCl, DNA was isolated with AMPure XP beads (beads:input of 84:100) and eluted in 30 μL 1 mM Tris-HCl pH 8.1. Samples were then subjected to 200-600 bp size selection using the BluePippin (Sage Science), prior to quantification of molarity using the Bioanalyzer (Agilent).

### CC-seq: NGS and data pipeline

Multiplexed library pools were sequenced on the Illumina MiSeq (Kit v3 −150 cycles) or Illumina NextSeq 500 (Kit v2 −75 cycles), with paired-end read lengths of 75 or 42 bp, respectively. Paired end reads that passed filter were aligned using bowtie2 (options: -X 1000 --no-discordant --very-sensitive --mp 5,1 --np 0), using MAPQ0 settings for yeast or MAPQ10 settings for human experiments, then SAM files processed by the custom-built Perl program termMapper that computes the coordinates of the protein-linked 5’-terminal nucleotide. The reference genomes used in this study are hg19 (human), and Cer3H4L2 (*S. cerevisiae*), which we generated by inclusion of the *his4*∷*LEU2* and *leu2*∷*hisG* loci into the Cer3 yeast genome build. Yeast data sets were filtered to exclude long terminal repeats, retrotransposons, telomeres, and the rDNA. Human datasets were filtered to remove known ultra-high signal regions (Hoffman et al., 2013; ENCODE, 2012) and repeat regions (ENCODE, 2012). All subsequent analyses were performed in R (Version 3.4.3) using RStudio (Version 1.1.383), unless indicated otherwise.

### CC-seq: Calibrated library generation

Calibration of Top2 CC-seq experiments was conducted to allow comparison of relative peak intensities in different yeast strains. This was achieved by spike-in of human DNA following the sonication stage of the protocol, which is a method that has been used to calibrate other sequencing methods (Hu et al., 2015; Grzybowski et al., 2015). S. cerevisiae and RPE-1 cells were exposed to etoposide and processed until just after sonication, exactly as according to the cell treatment and CC-seq protocols above. DNA concentration was quantified by Qubit, and then mixed at a molar ratio of human DNA:yeast DNA of 1:100. All subsequent stages of the protocol were identical, except read alignment, for which we used both hg19 and Cer3H4L2 builds successively. Cer3H4L2-aligned peak heights were corrected in each sample by multiplying by the reciprocal fraction of human reads in that sample.

### CC-seq: Fine-scale mapping

Fine-scale (nucleotide resolution) maps of Spo11/Top2 CCs were produced as simple histograms over a specified region (**Figures 1B, 2B, 3D, 4A, 5A, S1A, S3A, S5A and S5B**). Dark red and blue line heights indicate numbers of 5’-terminal nucleotides detected at that position on the Watson and Crick strands, in units of HpM. Where indicated in the figure caption, smoothed data are also plotted as pale red and blue polygons, in addition to the unsmoothed nucleotide resolution data. This smoothing was either applied using a sliding Hann window of the indicated width, or using a custom smoothing function (VarX), as indicated in the figure caption.

### CC-seq: Broad-scale mapping

Broad-scale maps of Top2 CCs were produced by binning nucleotide resolution data at 10 Kbp (**Figure 3B**) or 100 Kbp (**Figure S4D**) resolution. Binned data were either plotted directly (**Figure S4D**); or first scaled according to the estimated noise fraction (see below), smoothed with a 10-bin Hanning window, and median subtracted (**Figure 3B**).

### CC-seq: Estimation of the noise fraction in Human datasets

The noise fraction in each sample was estimated using an adaptation of the previously published NCIS method (Liang and Keleş, 2012). Briefly: the data for -VP16 and +VP16 samples were first binned at 10 Kbp resolution. Then the subpopulation of bins with the lowest TOP2 CC signal was identified in each sample. The average signal density of this subpopulation of bins, in each sample, was defined as the noise density (*d*_*-VP16*_ and *d*_*+VP16*_). Signal in the -VP16 sample was scaled by a normalisation factor equal to:

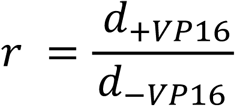

### CC-seq: Quantification of Spo11 and Top2 activity in defined genomic regions

Yeast Spo11 hotspots were the same as defined previously (Pan et al., 2011). Yeast intra/intergenomic regions were defined based on TSS and TTS coordinates reported on the Saccharomyces Genome Database (https://www.yeastgenome.org). Human chromatin compartments A and B were defined based on eigenvector analysis of previously published 100 kb resolution RPE-1 Hi-C data (Darrow et al., 2016) using the Juicer package (Lieberman-Aiden et al., 2009; Durand et al., 2016). Human “CTCF” and “TSS” regions were defined as 10 kb regions centred on the top quartile of expressed TSSs or the midpoint of top quartile CTCF-bound CTCF-binding motifs, with overlaps merged using GenomicFeatures∷reduce. The “Both” region was defined as the intersection of the “TSS” and “CTCF” regions using GenomicFeatures∷intersect. This region was then excluded from the individual “TSS” and “CTCF” regions. The “Neither” region was defined as regions of hg19 which overlapped neither “TSS”, nor “CTCF”, nor “Both”, using GenomicFeatures∷gaps. Nucleotide resolution data was tallied within defined regions, and is expressed as an aggregate (bar plots) in which the signal is corrected using the average signal in the “Neither” compartment, and/or as the distribution of region signal densities (box-and-whisker plot), as indicated in figure captions.

### CC-seq: Correlation of Watson/Crick cleavage positions

Nucleotide resolution human TOP2 data was thresholded at 0.01 HpM. Nucleotide resolution yeast Top2 and Spo11 data were not thresholded. Peak coordinates on the Crick strand were offset over the range −100 to +100, relative to Watson coordinates. After each offset, the data was filtered to include only sites with both Watson and Crick hits. The Pearson correlation (*r*) between the Watson and Crick signal intensities in these *n* sites was calculated. We also counted the fraction of reads found within these *n* sites, and normalised this number over the −100 to +100 bp range. Data are expressed as R values and/or normHpM values, as indicated in the figure captions.

### CC-seq: Aggregation of CC-seq data around loci of interest

Nucleotide resolution CC-seq data (line-plots) or binned CC-seq data (heatmaps) were aggregated within regions of specified size, centred on the loci of interest. The resulting sum total HpM was divided by numbers of loci to give a mean HpM per locus.

### CC-seq: Loci of interest used in this study

Yeast TSS and TTS coordinates were obtained from the Saccharomyces Genome Database https://www.yeastgenome.org. These were stratified based on associated gene length, expression level in vegetative SK1 (GSM907178, GSM907179, GSM907180), or expression level in meiotic SK1 (GSM907176, GSM907177; (Dominissini et al., 2012)). Human TSS and TTS coordinates were obtained from the UCSC hg19 knownGene annotation, and stratified based on associated gene length, or expression level (GSM1395252, GSM1395253, GSM1395254; (Ganem et al., 2014)). Occupied RPE-1 CTCF motifs were identified as follows: The FIMO tool (Grant et al., 2011) and the CTCF Position Weight Matrix (PWM) from the JASPAR database (Mathelier et al., 2014) were used to find all significant hg19 CTCF motifs (p < 1×10^−4^). These were filtered to include only those which overlapped positions of RPE-1 CTCF ChIP-seq peaks (GSM749673, GSM1022665; (ENCODE, 2012)), and stratified based on this ChIP-seq data. RPE-1 loop anchor-associated CTCF motifs were identified using the Juicer MotifFinder (Durand et al., 2016) with the RPE-1 WT Hi-C looplist (GSE71831 (Darrow et al., 2016)) and CTCF ChIP-seq BED file (GSM749673, GSM1022665; (ENCODE, 2012)) as input. Human TOP2 CC-seq peak coordinates were identified by thresholding (0.05 HpM) of pooled +VP16 data presented.

### CC-seq: Spatial correlation of yeast Top2 and Spo11 signals

Nucleotide resolution Top2/Spo11 CC-seq data was binned at 50 bp resolution and the Pearson correlation was calculated between the sum HpM for all bins at a given inter-bin distance.

### CC-seq: Quantifying single and double-strand Top2 CCs

Nucleotide resolution Human and Yeast Top2 CC-seq data were first thresholded at 0.05 and 1 HpM, respectively. SSB sites were defined as those sites without a cognate on the opposite strand at the expected offset of 3 bp. DSBs were defined as cognate sites with a 3 bp offset. SSB and DSB percentages are expressed as a percentage of the total number of sites (SSB + DSB). A randomisation experiment was conducted in order to estimate expected numbers of SSBs and DSBs within the sample under a random model where signal is distributed independently on each strand. To achieve this, the amplitudes of Top2 CCs (HpM) were shuffled amongst the positions in the nucleotide resolution datasets, prior to thresholding and offset analysis as described above.

### CC-seq: DNA sequence composition of single and double-strand Top2 CCs

Nucleotide resolution Human and Yeast Top2 CC-seq data were first thresholded at 0.05 and 1 HpM, respectively. The high frequency positions remaining were classified as part of a single or double-stranded Top2 CC based on absence or presence of a cognate cleavage at the expected W-C offset of 3 bp. For double-stranded Top2 CCs, a dyad axis coordinate was defined as the centrepoint between the Top2-linked nucleotides on the W and C strand (that is, the midpoint of the central two base pairs in the four base pair overhang). For single-stranded Top2 CCs, we used an imaginary dyad axis in the same relative position. DNA sequence was aggregated ±20 bp around these two classes of Top2 CCs. Statistical significance was determined using the one sample (goodness-of-fit) Chi-squared test, as described previously (Pommier et al., 1991).

### CC-seq -MethylC Analysis

A publically-available RPE-1 reduced representation bisulphite sequencing (RRBS) dataset (two replicates) was used to define coordinates of nucleotides immediately 3’-relative to 919,817 unmethylated or 341,136 methylated Cs, based on <10% or >90% methylation. Next we aggregated RPE-1 CC-seq (0.01 HpM thresholded) hits on these loci, and divided by the number of loci to give a density (barplot). Statistical significance was determined by the Kolmogorov-Smirnov test.

### Ideograms

Human chromosome ideograms were adapted from the open source ideogram package (https://eweitz.github.io/ideogram/).

## Supplementary Figure Legends

**Figure S1.**
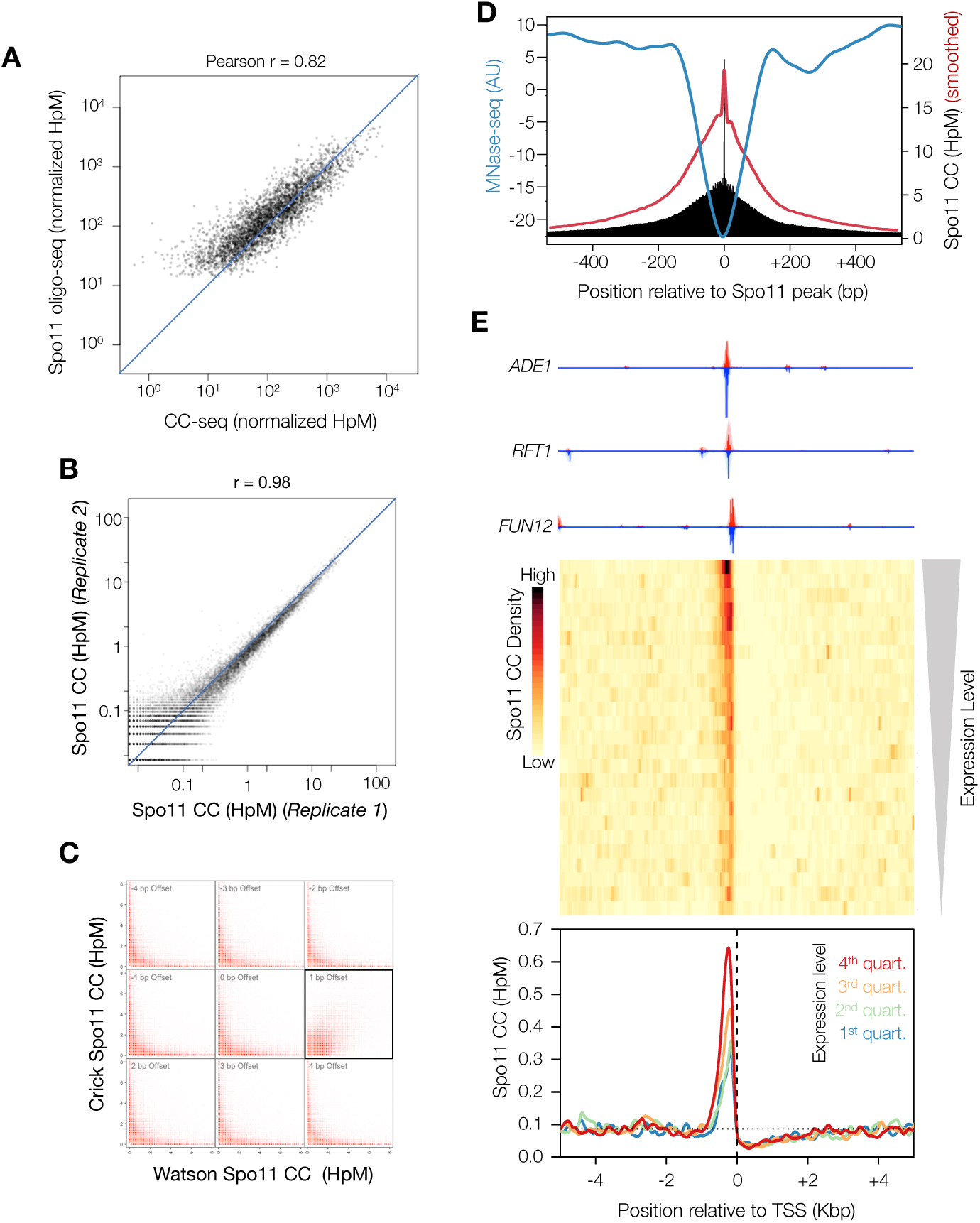
CC-seq maps covalent Spo11-linked DNA breaks in *S. cerevisiae* meiosis with nucleotide accuracy (related to Figure 1) **A)** Correlation of hotspot signals from CC-seq and oligo-seq. **B)** Correlation of 500 bp binned Spo11 maps from two representative replicates. **C)** Correlation of Spo11 cleavages on Watson and Crick strand when offset by 1 bp (boxed), relative to other offsets from −4 to +4 bp. **D)** Spo11 breaks mapped by CC-seq (raw=black, or smoothed=red) anticorrelate with nucleosome occupancy measured by MNase-seq (blue). **E)** Aggregation of Spo11 activity in a 10 Kbp window centred on orientated TSSs in *S. cerevisiae*. Three example TSSs are shown orientated in the 5’-3’ direction (top). A heatmap of all TSSs in the *S. cerevisiae* genome, with 25 rows stratified by gene expression level in SK1 cells (middle). The colour scale indicates average Spo11 break density. Motifs are also stratified into 4 quartiles of gene expression in meiotic SK1 cells, and the average distribution of Spo11 activity in each quartile plotted (bottom).

**Figure S2.**
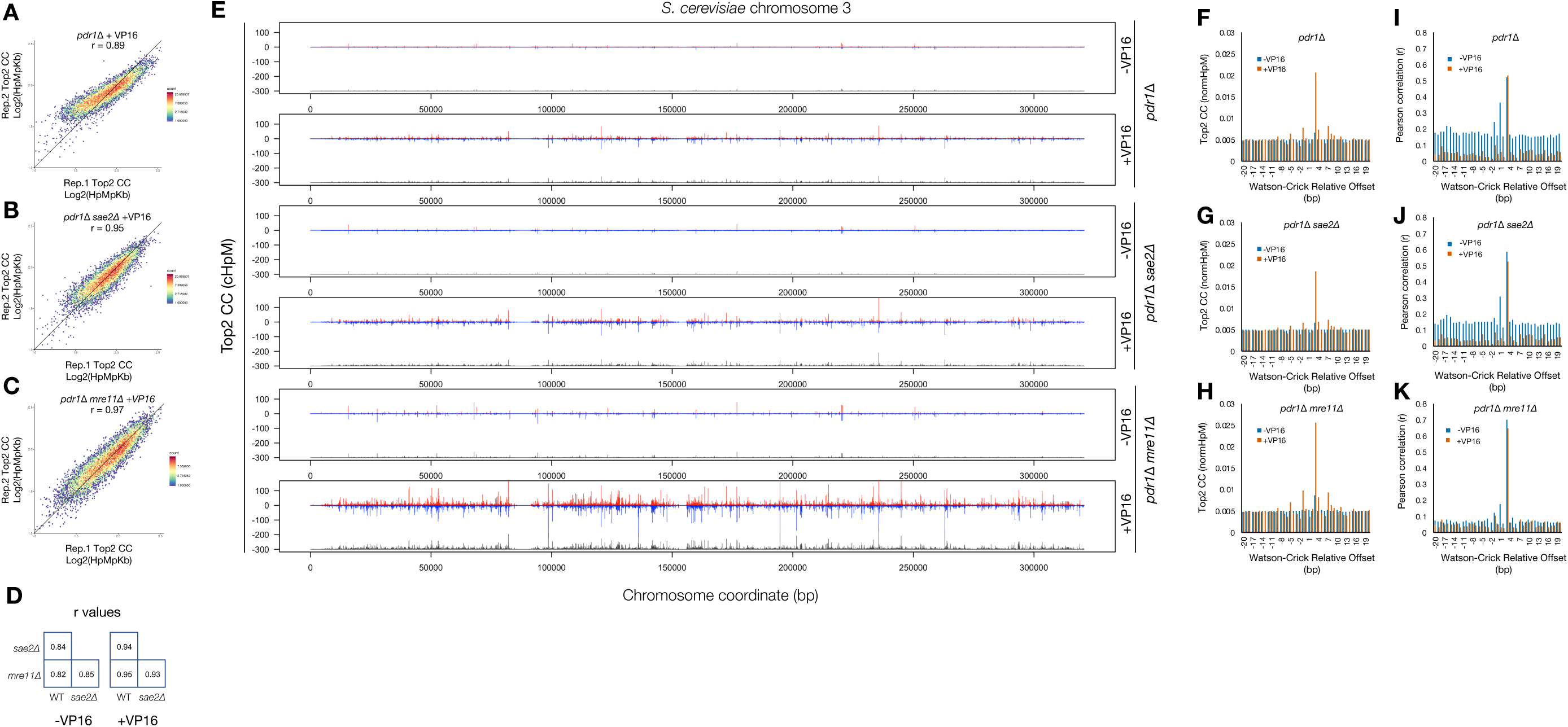
CC-seq maps covalent Top2-linked DNA breaks in *S. cerevisiae* cycling cells with nucleotide accuracy (related to Figure 2). **A-C)** Correlation of 1 Kbp binned Top2 CC maps from *pdr1*Δ (A), *pdr1*Δ*sae2*Δ (B) and *pdr1*Δ*mre11*Δ (C) cells treated with VP16. **D)** Pairwise Pearson correlation values for 100 bp binned Top2 CC maps from all assayed conditions. Each condition is a pool of two biological replicates. **E)** Nucleotide-resolution *S. cerevisiae* Top2 CC map of chromosome 3 for all assayed conditions. Each condition is a pool of two biological replicates. **F-H)** The normalised number of Top2 CCs retained in the *pdr1*Δ (F), *pdr1*Δ*sae2*Δ (G) and *pdr1*Δ*mre11*Δ (H) cells treated ±VP16, after filtering to include only sites offset by the given number of base pairs. All data were normalised over a −100 to +100 bp window. **I-K)** Pearson correlation (r) of Top2 CC-seq signal on Watson and Crick strands, offset by the indicated distance in *pdr1*Δ (I), *pdr1*Δ*sae2*Δ (J) and *pdr1*Δ*mre11*Δ (K) cells treated ±VP16.

**Figure S3.**
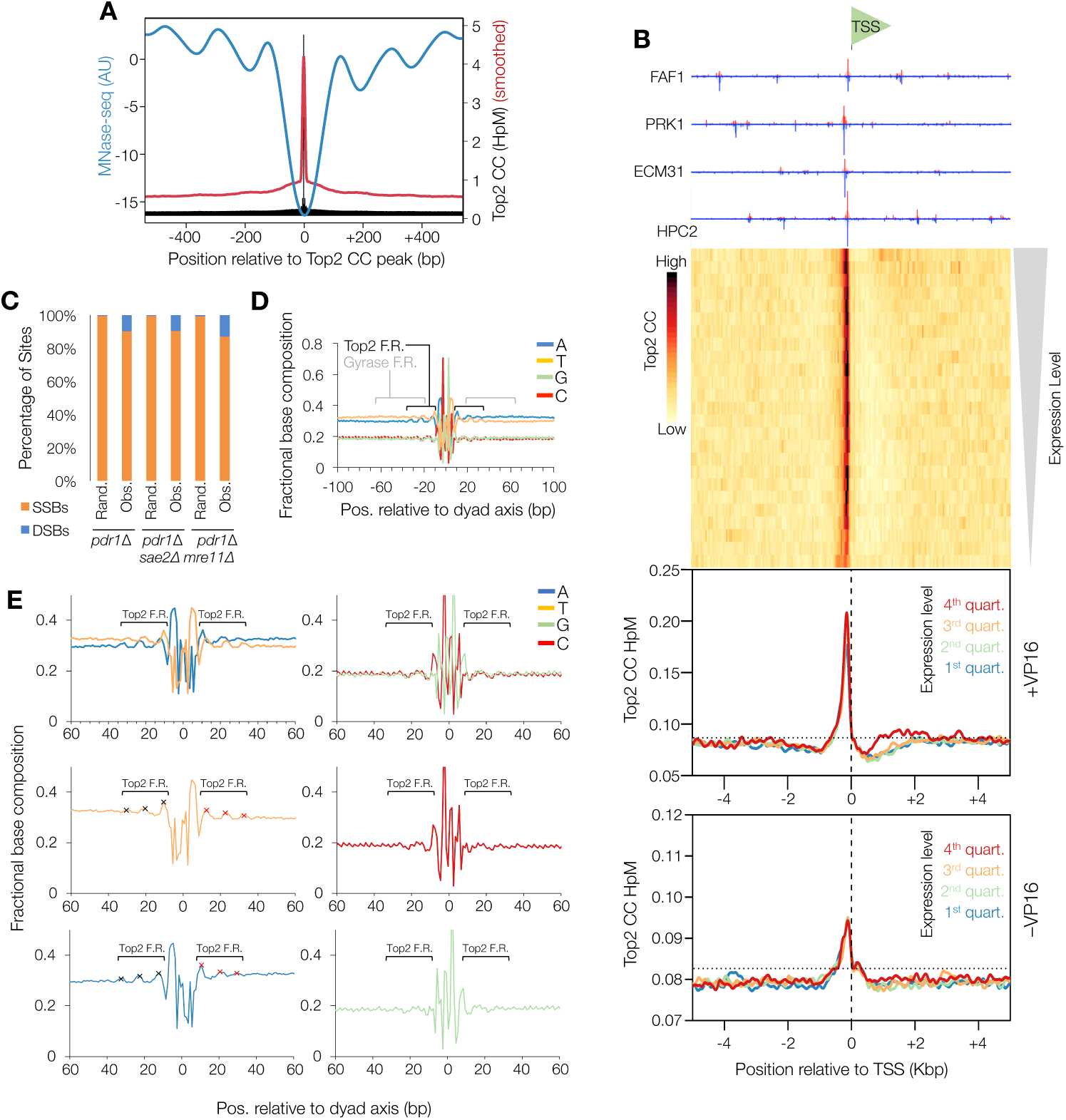
Top2-linked DNA breaks in *S. cerevisiae* are not correlated with gene expression, anticorrelate with nucleosomes, and have biased nucleotide skews indicative of bent DNA (related to Figure 2) **A)** Top2 CC mapped by CC-seq (raw=black, or smoothed=red) anticorrelate with nucleosome occupancy measured by MNase-seq (blue). **B)** Aggregation of Top2 CCs in a 10 Kbp window centred on orientated transcription start sites (TSS) in *pdr1*Δ *S. cerevisiae*. Four example TSSs are shown orientated in the 5’-3’ direction (top). Heatmap of all TSSs in the *S. cerevisiae* genome, with 25 rows stratified by gene expression level in vegetative growth (middle). Colour scale indicates average Top2 CC density. Motifs are also stratified into four quartiles of gene expression, and the average distribution of Top2 CCs in each quartile plotted for untreated and +VP16 conditions (bottom). **C)** The percentage of strong sites that are SSBs and DSBs in etoposide-treated *pdr1*Δ, *pdr1*Δ*sae2*Δ and *pdr1*Δ*mre11*Δ *S. cerevisiae*. Sites were thresholded at 1 HpM prior to sorting into DSB or SSB classes based on presence or absence of a 3 bp offset cognate (Obs.). As a control, the amplitudes of Top2 CCs (HpM) were randomised amongst the positions in the nucleotide resolution datasets, prior to thresholding and offset analysis as described above (Rand.) **D)** Average nucleotide composition over a 200 bp window centred on the DSB dyad axis, in etoposide-treated *pdr1*Δ *S. cerevisiae*. The position of the flanking regions (F.R.) identified in Gyrase mapping experiments, and the flanking regions observed here for Top2 are indicated in blue and black, respectively. **E)** Average nucleotide composition over a 120 bp window centred on the DSB dyad axis, in etoposide-treated *pdr1*Δ *S. cerevisiae*. The signal pattern is separated into A+T and G+C (top left and right), and into individual A, T, C and G plots (middle left, bottom left, middle right, and bottom right, respectively). The position of the flanking regions (F.R.) observed here for Top2 are indicated in black, and their ∼10.5 bp periodicity is highlighted with black and red crosses.

**Figure S4.**
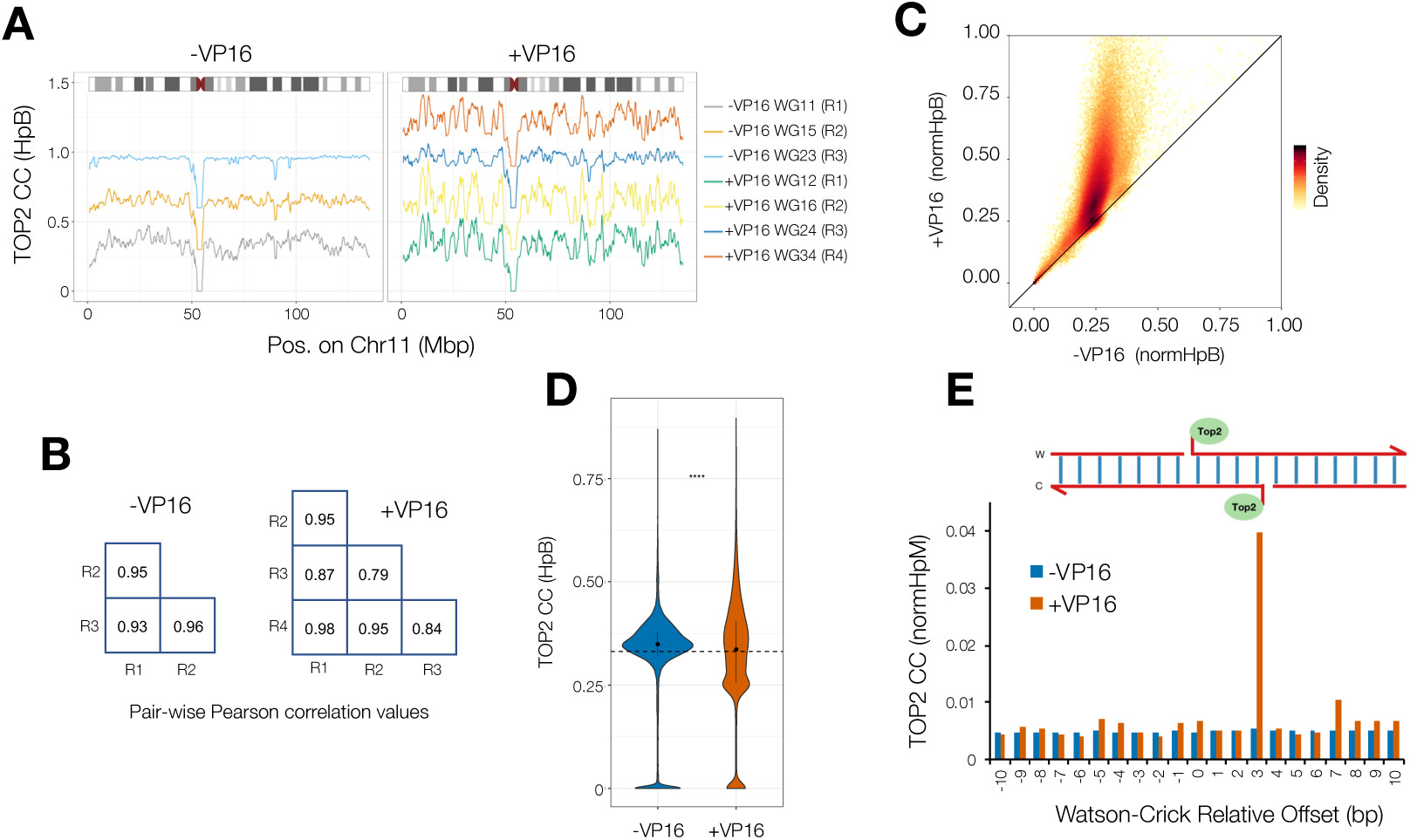
CC-seq maps of TOP2-linked DNA breaks in Human cells are enriched by etoposide, and show high reproducibility and nucleotide accuracy (related to Figure 3) **A)** Broad-scale *H. sapiens* TOP2 CC-seq maps in individual biological replicates of RPE-1 cells ±VP16. Raw data were binned at 100 Kbp prior to plotting. Each plot is offset on the y-axis by +0.3 HpB. **B)** Replicate-to-replicate Pearson correlation values (r) for 10 Kbp binned TOP2 CC-seq maps of RPE-1 cells ±VP16. **C)** Scatter plot of -VP16 and +VP16 TOP2 CC-seq maps binned at 10 Kbp resolution. Data were first scaled according to the estimated noise fraction (**Methods**), and are presented in a hexagonal-binned format, where the density of overplotting is indicated by the colour scale. **D)** Violin plots of TOP2 CC-seq maps ±VP16 binned at 100 Kbp resolution. The inner black bar, black dot, and dotted horizontal line indicate the interquartile range, median, and expected mean Top2 CC density based on random distribution. **E)** The normalised number of TOP2 CCs retained in the CC-seq maps in RPE-1 cells ±VP16 after filtering to include only sites offset by the given number of base pairs. Data were normalised over a −100 to +100 bp window.

**Figure S5.**
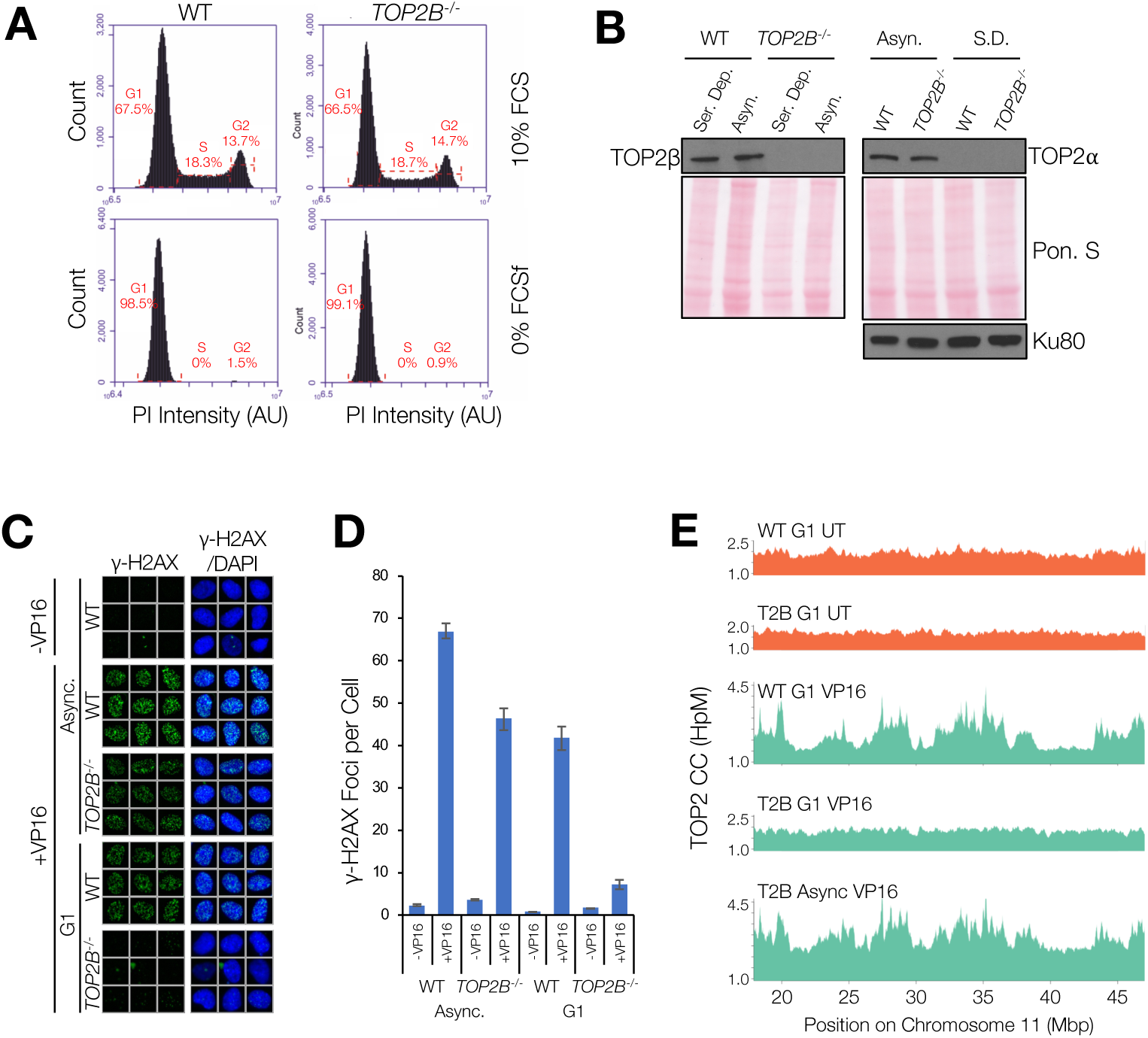
CC-seq signal is TOP2-dependent. **A)** DNA content histograms of wild type (WT) and *TOP2B*^−/−^ RPE-1 cells under asynchronous (10% FCS) and serum-deprived (0% FCS) conditions, as measured by FACS following propidium iodide (PI) staining. G1, S and G2 populations are clearly present under asynchronous growing conditions. A strong G1 arrest is observed in serum deprived conditions. Percentages of cells in each of the indicated regions (red dotted brackets) are given. **B)** Western blots demonstrating the absence of TOP2β protein in serum deprived and asynchronous *TOP2B*^−/−^ RPE-1 cells (left), and the absence of TOP2α in serum deprived wild type and *TOP2B*^−/−^ RPE-1 cells (right). Ponceau S total protein loading is presented (Pon. S) for the left and right panels, and additionally a Ku80 loading control is included for the right panel. **C)** Immunofluorescence experiment demonstrating induction of γ-H2AX foci (green) in asynchronous (Async.) and serum-deprived (Ser. Dep.) wild type (WT) and *TOP2B*^−/−^ RPE-1 cells, all co-stained with DAPI (blue). Galleries of nine cells per condition were chosen randomly using Olympus ScanR Analysis software. **D)** Quantification of (C). Numbers of γ-H2AX foci per cell were counted automatically using Olympus ScanR Analysis software. The mean ±SEM is reported for n= 3 biological replicate experiments. **E)** Broad-scale *H. sapiens* TOP2 CC-seq maps in asynchronous and serum-deprived wild type (WT) and *TOP2B*^−/−^ RPE-1 cells -VP16 (orange) and +VP16 (green). Raw hits on Watson and Crick strands were summed and smoothed according to local signal density (Fsize=501).

**Figure S6.**
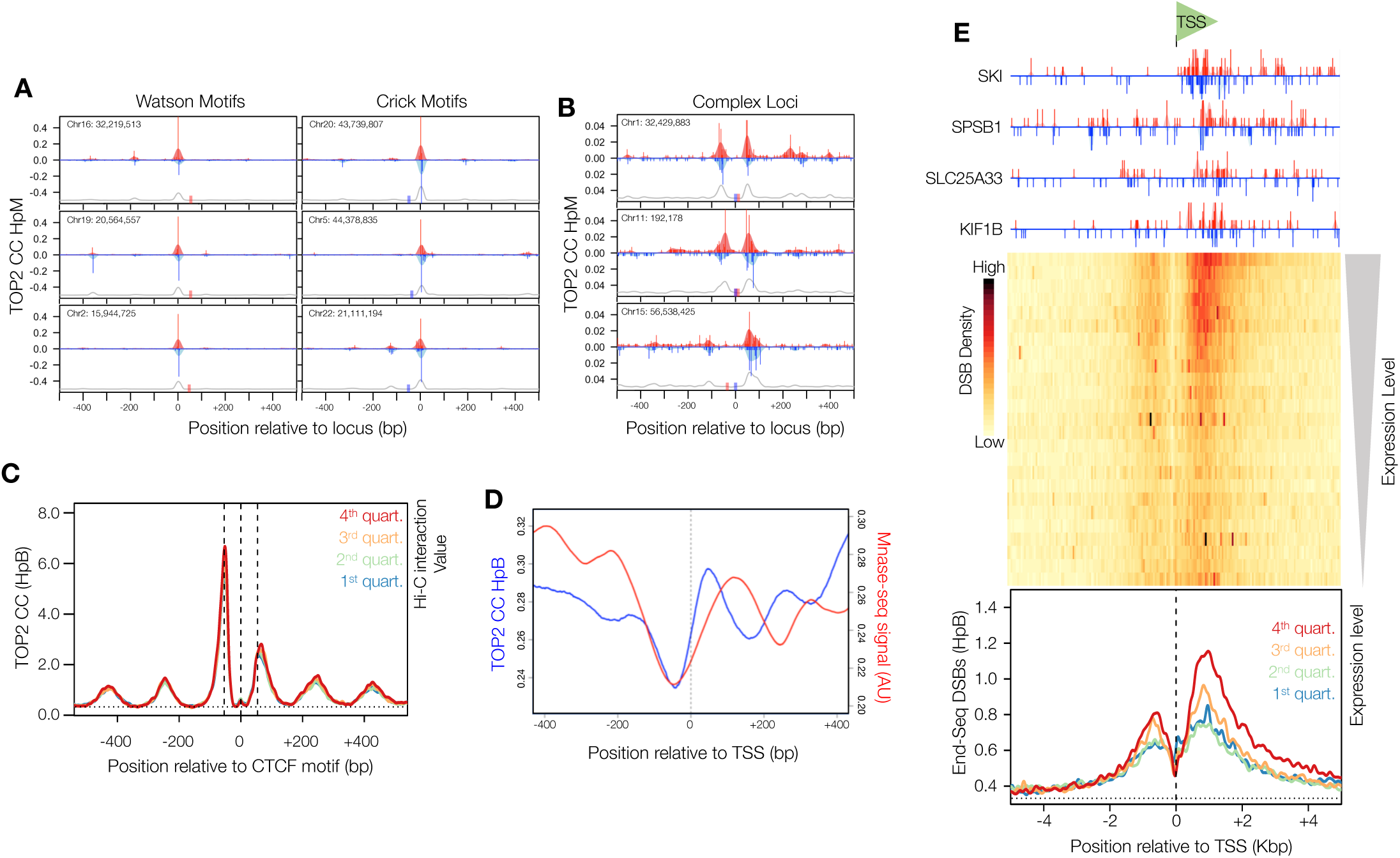
TOP2 CC-seq signal enrichment around CTCF and TSS sites, compared with END-seq DSB signal at TSSs. **A)** Fine-scale TOP2 CC-seq maps of *H. sapiens* CTCF-proximal loci in RPE-1 cells +VP16. Red and blue traces indicate TOP2-linked 5’ DNA termini on the Watson and Crick strands, respectively. Pale shaded areas are the same data smoothed with a sliding 11 bp Hanning window. Red and Blue rectangles indicate the positions of CTCF motifs on the Watson and Crick strands respectively. The grey line indicates Hanning-smoothed sum of Watson and Crick TOP2 CCs. **B)** Fine-scale mapping of TOP2 CCs surrounding three complex CTCF loci, processed as in (A). **C)** Aggregation of TOP2 CCs in a 1 Kbp window centred on the subset of orientated CTCF motifs that can be assigned to a chromatin loop anchor in human RPE-1 cells (Darrow et al., 2016). Motifs are stratified into 4 quartiles of loop anchor interaction strength, and the average TOP2 CC distribution in each quartile plotted. **D)** Fine-scale aggregation of TOP2 CCs (red) in a 800 bp window centred on TSSs in human RPE-1 cells, showing anticorrelation with aggregated MNase-seq signal (blue). **E)** Aggregation of END-seq mapped DSBs in a 10 Kbp window centred on orientated TSSs in human MCF7 cells. Four example TSSs are shown orientated in the 5’-3’ direction (top). A heatmap of all TSSs in the human genome, with 25 rows stratified by flanking gene expression level in MCF7 cells (middle). The colour scale indicates average END-seq DSB density. TSSs are also stratified into 4 quartiles based on strength of flanking gene expression, and the average END-seq DSB distribution in each quartile plotted (bottom).

**Figure S7.**
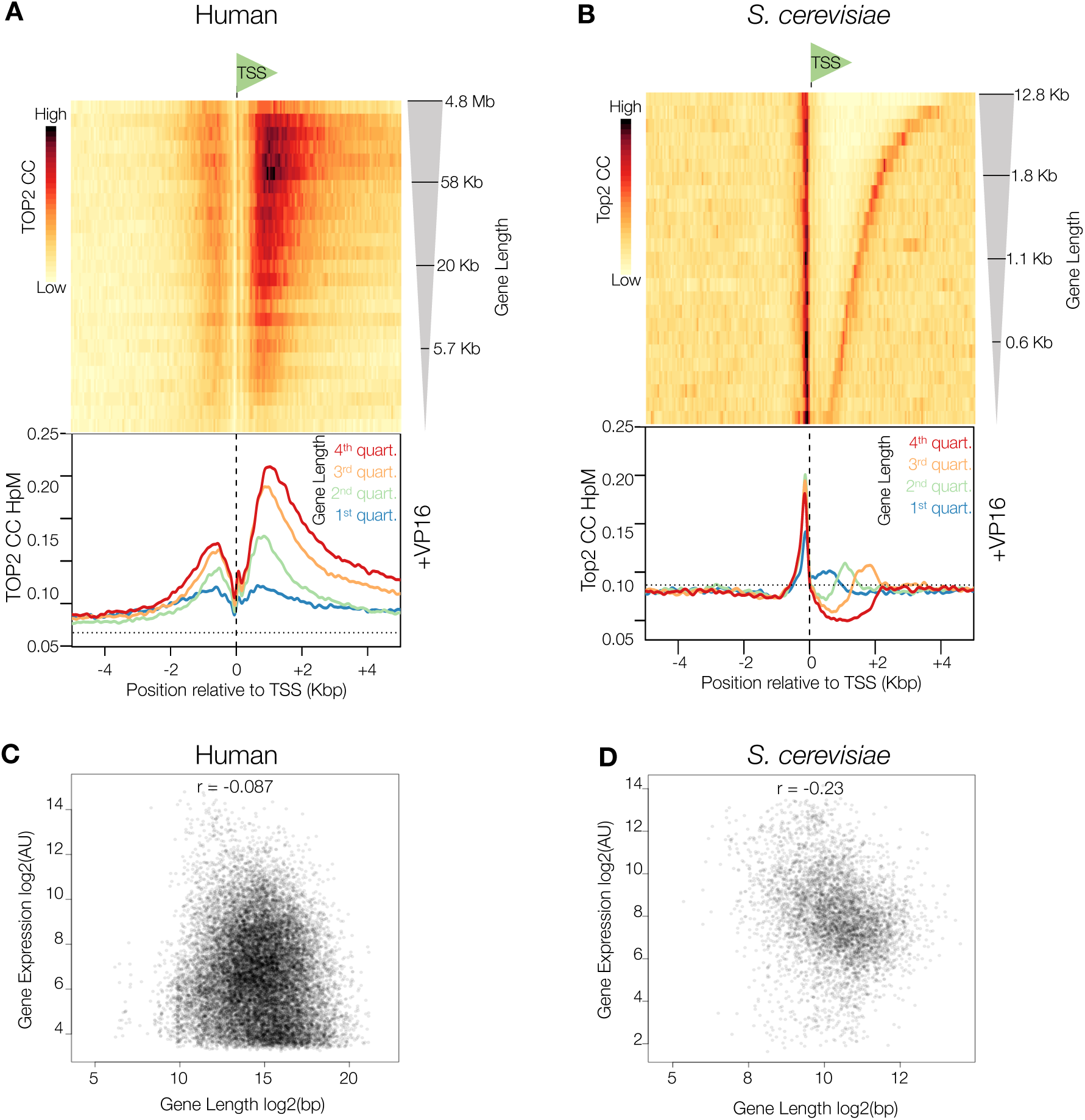
TSS-proximal TOP2-linked DNA breaks in humans are correlated with gene length, independently from gene expression level (related to Figure 5); TSS-proximal Top2-linked DNA breaks in yeast are not strongly correlated with gene length, (related to Figure 2) **A)** Aggregation of TOP2 CCs in a 10 Kbp window centred on orientated TSSs in human RPE-1 cells. A heatmap of all TSSs in the human genome, with 25 rows stratified by gene length (top). The colour scale indicates average TOP2 CC density. Motifs are also stratified into 4 quartiles of gene length, and the average TOP2 CC distribution in each quartile plotted (bottom). **B)** Aggregation of Top2 CCs in a 10 Kbp window centred on orientated transcription start sites (TSS) in *pdr1*Δ *S. cerevisiae*. Heatmap of all TSSs in the *S. cerevisiae* genome, with 25 rows stratified by gene length level (top). Colour scale indicates average Top2 CC density. Motifs are also stratified into four quartiles of gene length, and the average distribution of Top2 CCs in each quartile plotted (bottom). **C)** Scatter plot of human gene length and RPE-1 gene expression, showing no correlation. **D)** Scatter plot of *S. cerevisiae* gene length and gene expression during vegetative growth, showing little correlation.

## Supplementary Tables

**Table S1.**
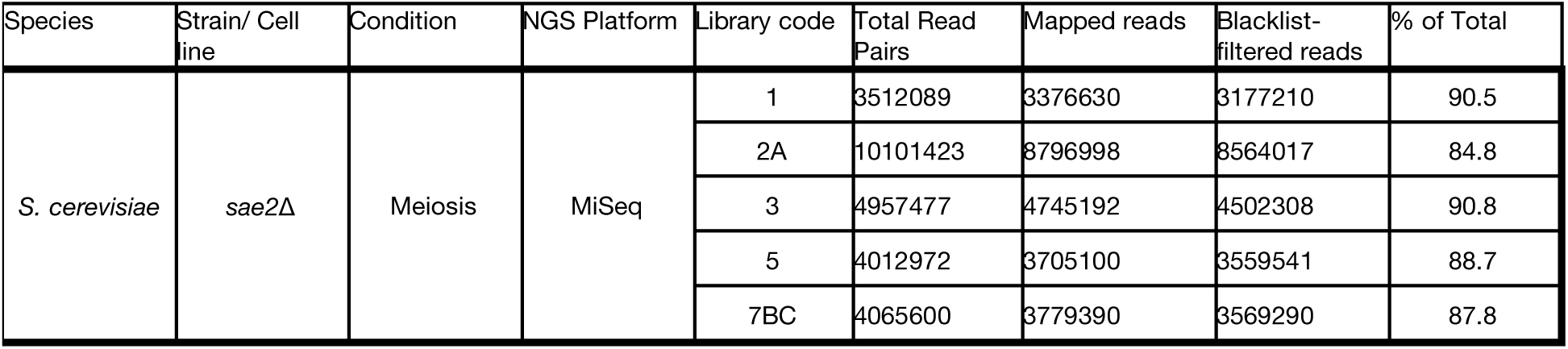
DNA libraries used for mapping Spo11 activity. The number of reads remaining after each stage of the data pipeline are indicated.

**Table S2.** DNA libraries used for mapping Top2 activity. The number of reads remaining after each stage of the data pipeline are indicated.

| Species | Strain/ Cell line | Condition | NGS Platform | Library code | Total Read Pairs | Mapped reads | De-duplicated reads | Blacklist-filtered reads | % of Total | Pooled Reads (Millions) |
| --- | --- | --- | --- | --- | --- | --- | --- | --- | --- | --- |
| S. cerevisiae | pdr1Δ | -VP16 | NextSeq 500 | RA1 | 3741251 | 3542955 | 3179049 | 2746294 | 73.4 | 3.50 |
|  |  |  | MiSeq | RA8 | 1118483 | 879107 | 842290 | 751850 | 67.2 |  |
| S. cerevisiae | pdr1Δ | +VP16 | NextSeq 500 | RA2 | 3096257 | 3053356 | 2679303 | 2234941 | 72.2 | 3.93 |
|  |  |  | MiSeq | RA9 | 2262958 | 2071231 | 1958341 | 1690075 | 74.7 |  |
| S. cerevisiae | pdr1Δ sae2Δ | -VP16 | NextSeq 500 | RA4 | 3634284 | 3552818 | 3215523 | 2745668 | 75.5 | 3.79 |
|  |  |  | MiSeq | RA10 | 1420662 | 1214067 | 1181399 | 1046827 | 73.7 |  |
| S. cerevisiae | pdr1Δ sae2Δ | +VP16 | NextSeq 500 | RA5 | 2529368 | 2476714 | 1386551 | 1097282 | 43.4 | 3.54 |
|  |  |  | MiSeq | RA11 | 3226626 | 3063388 | 2824882 | 2444486 | 75.8 |  |
| S. cerevisiae | pdr1Δ mre11Δ | -VP16 | NextSeq 500 | RA6 | 5345587 | 5205411 | 4288662 | 3646842 | 68.2 | 4.76 |
|  |  |  | MiSeq | RA12 | 1519020 | 1326435 | 1280137 | 1109310 | 73.0 |  |
| S. cerevisiae | pdr1Δ mre11Δ | +VP16 | NextSeq 500 | RA7 | 3382546 | 3357168 | 2921333 | 2526342 | 74.7 | 5.54 |
|  |  |  | MiSeq | RA13 | 4133347 | 4032996 | 3540366 | 3010542 | 72.8 |  |
| Human | RPE-1 WT | -VP16 | NextSeq 500 | WG11 | 125203096 | 109503841 | 68845619 | 68639594 | 54.8 | 185.36 |
|  |  |  |  | WG15 | 125328792 | 110432447 | 46588296 | 46449378 | 37.1 |  |
|  |  |  |  | WG23 | 89154181 | 77728045 | 70484665 | 70269334 | 78.8 |  |
| Human | RPE-1 WT | +VP16 | NextSeq 500 | WG12 | 110910655 | 95546007 | 55848200 | 55681939 | 50.2 | 276.97 |
|  |  |  |  | WG16 | 102804692 | 91872706 | 54312084 | 54164650 | 52.7 |  |
|  |  |  |  | WG24 | 130816235 | 114086421 | 101162102 | 100857557 | 77.1 |  |
|  |  |  |  | WG34 | 91829516 | 79751337 | 66458538 | 66270349 | 72.2 |  |
| Human | RPE-1 WT | -VP16 G1 | MiSeq | WG13 | 2601642 | 2292863 | 2007177 | 1991088 | 76.5 | 4.27 |
|  |  |  |  | WG21 | 2610523 | 2298289 | 2295625 | 2277124 | 87.2 |  |
| Human | RPE-1 WT | +VP16 G1 | MiSeq | WG14 | 3075620 | 2771819 | 2491674 | 2475350 | 80.5 | 5.23 |
|  |  |  |  | WG22 | 3139164 | 2779809 | 2772349 | 2751281 | 87.6 |  |
| Human | RPE-1 WT | -VP16 Async | MiSeq | WG15b* | 3207507 | 2866283 | 2782876 | 2760652 | 86.1 | 5.35 |
|  |  |  |  | WG23 | 2959289 | 2616954 | 2608670 | 2586523 | 87.4 |  |
| Human | RPE-1 WT | +VP16 Async | MiSeq | WG16b* | 3018163 | 2721770 | 2673273 | 2656580 | 88.0 | 6.30 |
|  |  |  |  | WG24 | 4165048 | 3682708 | 3668602 | 3639324 | 87.4 |  |
| Human | RPE-1 TOP2B- | -VP16 G1 | MiSeq | WG17 | 2978044 | 2624262 | 2558332 | 2536752 | 85.2 | 4.69 |
|  |  |  |  | WG25 | 2460949 | 2181002 | 2173777 | 2155101 | 87.6 |  |
| Human | RPE-1 TOP2B- | +VP16 G1 | MiSeq | WG18 | 2927745 | 2588870 | 2557532 | 2537085 | 86.7 | 5.10 |
|  |  |  |  | WG26 | 2922624 | 2596547 | 2587152 | 2565454 | 87.8 |  |
| Human | RPE-1 TOP2B- | -VP16 Async | MiSeq | WG19 | 3713068 | 3278164 | 3188049 | 3162895 | 85.2 | 7.04 |
|  |  |  |  | WG27 | 4413242 | 3943962 | 3910035 | 3877611 | 87.9 |  |
| Human | RPE-1 TOP2B- | +VP16 Async | MiSeq | WG20 | 3426100 | 3038601 | 2978047 | 2956076 | 86.3 | 8.13 |
|  |  |  |  | WG28 | 5878342 | 5251089 | 5211027 | 5169010 | 87.9 |  |

**Table S3.** *S. cerevisiae* strains used in this study. Strains MJ315 and M319 were used for Spo11 mapping; strains MJ429, MJ475 and MJ551 were used for Top2 mapping.

| Strain | Number | Genotype |
| --- | --- | --- |
| *sae2Δ* | MJ315 | SK1: *ho::LYS2'*, *lys2'*, *ura3'*, *arg4-nsp'*, *leu2::hisG'*, *his4X::LEU2'*, *nuc1::LEU2'*, *sae2Δ::KanMX6'* |
| *sae2Δ spo11-Y135F* | MJ319 | SK1: *ho::LYS2'*, *lys2'*, *ura3'*, *arg4-nsp'*, *leu2::hisG'*, *his4X::LEU2'*, *nuc1::LEU2'*, *spo11(Y135F)-HA3His6::KanMX4'*, *sae2Δ::KanMX6'* |
| *pdr1Δ* | MJ429 | BY4741: *ura3Δ0*, *leu2Δ0*, *his3Δ1*, *met15Δ0*, *pdr1Δ::PDR1-DBD-CYC8::LEU2* |
| *pdr1Δ sae2Δ* | MJ475 | BY4741: *ura3Δ0*, *leu2Δ0*, *his3Δ1*, *met15Δ0*, *pdr1Δ::PDR1-DBD-CYC8::LEU2*, *sae2Δ::KanMX6* |
| *pdr1Δ mre11Δ* | MJ551 | BY4741: *ura3Δ0*, *leu2Δ0*, *his3Δ1*, *met15Δ0*, *pdr1Δ::PDR1-DBD-CYC8::LEU2*, *mre11Δ::KanMX4* |

**Table S4.** Human cell lines used in this study.

| Parent cell line | Clone number | Genotype |
| --- | --- | --- |
| hTERT RPE-1 | WT | WT |
| hTERT RPE-1 | T2B/1 | *TOP2B* |

**Table S5.**
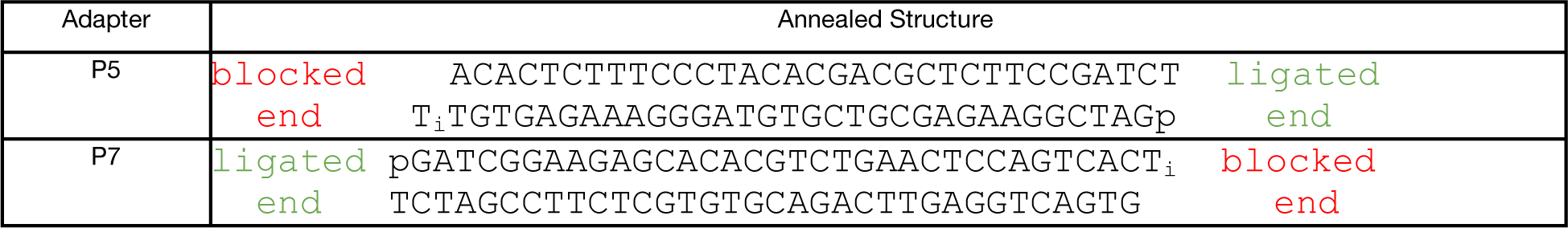
The annealed structures of the custom P5 and P7 adapters. “T_i_” indicates an inverted dT, linked via a 3’-3’ phosphodiester. This inhibits unwanted ligation at this end of the adapter. Also note that the 5’-terminal moieties of each adapter (A for P5, G for P7) are nucleosides (3’-OH), which further inhibits ligation at this end

**Table S6.** Publically available datasets used in this study.

| Species/ Cell line | Dataset Type | Accession(s) | Reference |
| --- | --- | --- | --- |
| *S. cerevisiae* / SK1 (meiotic) | Gene expression microarray | GSM907178, GSM90719, GSM907180 | (Dominissini et al., 2012) |
| *S. cerevisiae* / SK1 (vegetative) | Gene expression microarray | GSM907176, GSM907177 | (Dominissini et al., 2012) |
| *H. sapiens* / RPE-1 | Gene expression microarray | GSM1395252, GSM1395253, GSM1395254 | (Ganem et al., 2014) |
| *H. Sapiens* / MCF7 | Gene expression microarray | GSM1141244 | (Nelson et al., 2013) |
| *H. sapiens* / RPE-1 | Hi-C | GSE71831 | (Darrow et al., 2016) |
| *H. sapiens* / RPE-1 | bTMP-seq | GSM1062645, GSM1062646 | (Naughton et al., 2013) |
| *H. sapiens* / MCF7 | END-seq | GSM2635568 | (Canela et al., 2017) |
| *H. sapiens* / RPE-1 | CTCF ChIP-seq | GSM749673, GSM1022665 | (ENCODE, 2012) |
| *H. sapiens* / RPE-1 | H3K4Me3 ChIP-seq | GSM945271, GSM945271 | (ENCODE, 2012) |
| *H. sapiens* / RPE-1 | H3K27Ac ChIP-seq | GSM733771, GSM733718, GSM733755, GSM733691, GSM733656, GSM733674, GSM733646 | (ENCODE, 2012) |
| *H. sapiens* / RPE-1 | Methyl-RRBS | GSM683773, GSM683905 | (ENCODE, 2012) |
| *H. sapiens* / K562 | MNase-seq | GSM920557 | (ENCODE, 2012) |

